# Shared PRAME Epitopes are T-Cell Targets in NUT Carcinoma

**DOI:** 10.1101/2025.03.07.642090

**Authors:** Jeffrey L. Jensen, Sara K. Peterson, Maria Sambade, Jessica R. Alley, Shawn Yu, Tomoaki Kinjo, Sarah N. Bennett, Steven P. Vensko, Mitra Shabrang, Johnathan D. DeBetta, Julie K. Geyer, Brandon A. Price, Kwangok P. Nickel, Randall J. Kimple, Rishi S. Kotecha, Laura E. Herring, Ian J. Davis, Jeremy R. Wang, Christopher A. French, Brian Kuhlman, Jared Weiss, Alex Rubinsteyn, Benjamin G. Vincent

**Affiliations:** Division of Medical Oncology, Department of Medicine, The University of North Carolina at Chapel Hill, Chapel Hill, North Carolina; Lineberger Comprehensive Cancer Center, Chapel Hill, North Carolina; Department of Genetics, Curriculum in Bioinformatics and Computational Biology, The University of North Carolina at Chapel Hill, Chapel Hill, North Carolina; Department of Biochemistry and Biophysics, University of North Carolina at Chapel Hill, Chapel Hill, North Carolina; Department of Microbiology and Immunology, The University of North Carolina at Chapel Hill, Chapel Hill, North Carolina; Curriculum in Toxicology and Environmental Medicine, School of Medicine, University of North Carolina at Chapel Hill, Chapel Hill, North Carolina; Department of Biochemistry and Molecular Biology, John Hopkins Bloomberg School of Public Health, Baltimore, Maryland; Department of Pathology and Laboratory Medicine, University of North Carolina at Chapel Hill, Chapel Hill, North Carolina; Tempus AI, Inc., Chicago Illinois; Department of Human Oncology, University of Wisconsin School of Medicine and Public Health, University of Wisconsin, Madison, WI; University of Wisconsin Carbone Cancer Center, University of Wisconsin School of Medicine and Public Health, University of Wisconsin, Madison, WI; Department of Clinical Haematology, Oncology, Blood and Marrow Transplantation, Perth Children’s Hospital, Perth, Australia; Leukaemia Translational Research Laboratory, WA Kids Cancer Centre, The Kids Research Institute Australia, University of Western Australia, Perth, Australia; Curtin Medical School, Curtin University, Perth, Australia; Division of Hematology-Oncology, Department of Pediatrics, The University of North Carolina at Chapel Hill, Chapel Hill, North Carolina; Department of Pathology, Brigham and Women’s Hospital, Harvard Medical School, Boston, Massachusetts; Division of Hematology, Department of Medicine, The University of North Carolina at Chapel Hill, Chapel Hill, North Carolina

**Keywords:** NUT carcinoma, PRAME, T-Cell Receptor Bispecifics, Cancer-specific Antigens, brenetafusp, IMC-F106C, BRD4::NUTM1, BRD3::NUTM1, NSD3::NUTM1, anzutresgene autoleucel

## Abstract

**Background:** NUT carcinoma is a rare but highly lethal solid tumor without an effective standard of care. NUT carcinoma is caused by bromodomain-containing *NUTM1* fusion oncogenes, most commonly *BRD4::NUTM1*. BRD4::NUTM1 recruits p300 to acetylate H3K27 forming expansive stretches of hyperacetylated chromatin called “megadomains” with the overexpression of corresponding oncogenes, including *MYC*. We hypothesized that transcriptional dysregulation caused by BRD4::NUTM1 would lead to the generation of cancer-specific antigens that could be therapeutically actionable.

**Methods:** We integrated genomics, computational antigen prediction software, targeted immunopeptidomics using single- and double-labeled peptide standards, and gain/loss-of-function genetic experiments on a panel of cell lines (N=5), a patient derived xenograft, a tissue microarray (N=77), and patient samples from the Tempus AI Sequencing Database harboring evidence of *NUTM1* fusions (N=165). We created an αPRAME_425_ T-cell receptor x SP34 αCD3 bispecific molecule modeled after brenetafusp, an αPRAME_425_ T-cell receptor bispecific T-cell engager, as well as αPRAME_425_ TCR T-cells based on anzutresgene autoleucel and we applied these products to NUT carcinoma cells *in vitro*.

**Results:** We identified *PRAME* as the most commonly expressed cancer/testis antigen in patient samples harboring the three canonical NUT carcinoma fusions (*BRD4::NUTM1*, *BRD3::NUTM1*, and *NSD3::NUTM1*). Additionally, 56% (43/77) of NUT carcinoma tissue microarray samples stained positive for PRAME. *BRD4::NUTM1* expression in HEK 293T cells enhanced PRAME levels and *BRD4::NUTM1* knockout in NUT carcinoma cells reduced PRAME levels. Immunopeptidomics detected more PRAME-derived HLA ligands (N=9) than all other cancer/testis antigens combined (N=5). Targeted mass spectrometry detected the HLA-A*02:01/SLLQHLIGL (PRAME_425_) epitope in 100% (4/4) of HLA-A*02+, PRAME+ NUT carcinoma samples at higher levels (>0.01 fM) than HLA-A*02:01/RLDQLLRHV (PRAME_312_) or HLA-A*02:01/YLHARLREL (PRAME_462_). The αPRAME_425_ T-cell receptor x SP34 αCD3 bispecific molecule and αPRAME_425_ TCR T-cells each exhibited potent, T-cell mediated cytotoxicity against *PRAME*+ NUT carcinoma cells.

**Conclusions:** *PRAME* is highly and frequently expressed in NUT carcinoma and the most common oncoprotein causing NUT carcinoma, BRD4::NUTM1, contributes to these high PRAME levels. PRAME epitopes presented by HLA Class I are a previously unrecognized therapeutic vulnerability for NUT carcinoma that warrant clinical trials testing PRAME targeted immunotherapies in this neglected patient population.

**What is already known on this topic:** NUT carcinoma is a devastating malignancy that is recalcitrant to cytotoxic chemotherapy, T-cell checkpoint blockade, and targeted therapies in the form of bromodomain inhibitors.

**What this study adds:** NUT carcinoma tumors are high in the cancer/testis gene *PRAME*. The oncogene most commonly causing NUT carcinoma, *BRD4::NUTM1*, contributes to these high levels. NUT carcinoma cells present PRAME epitopes on HLA Class I molecules and are susceptible to PRAME-directed, T-cell mediated cytotoxicity.

**How this study might affect research, practice or policy:** Our results argue for phase I/II clinical trials testing PRAME immunotherapies like brenetafusp or anzutresgene autoleucel in *PRAME*+ NUT carcinoma patients.

**Graphical Abstract:** 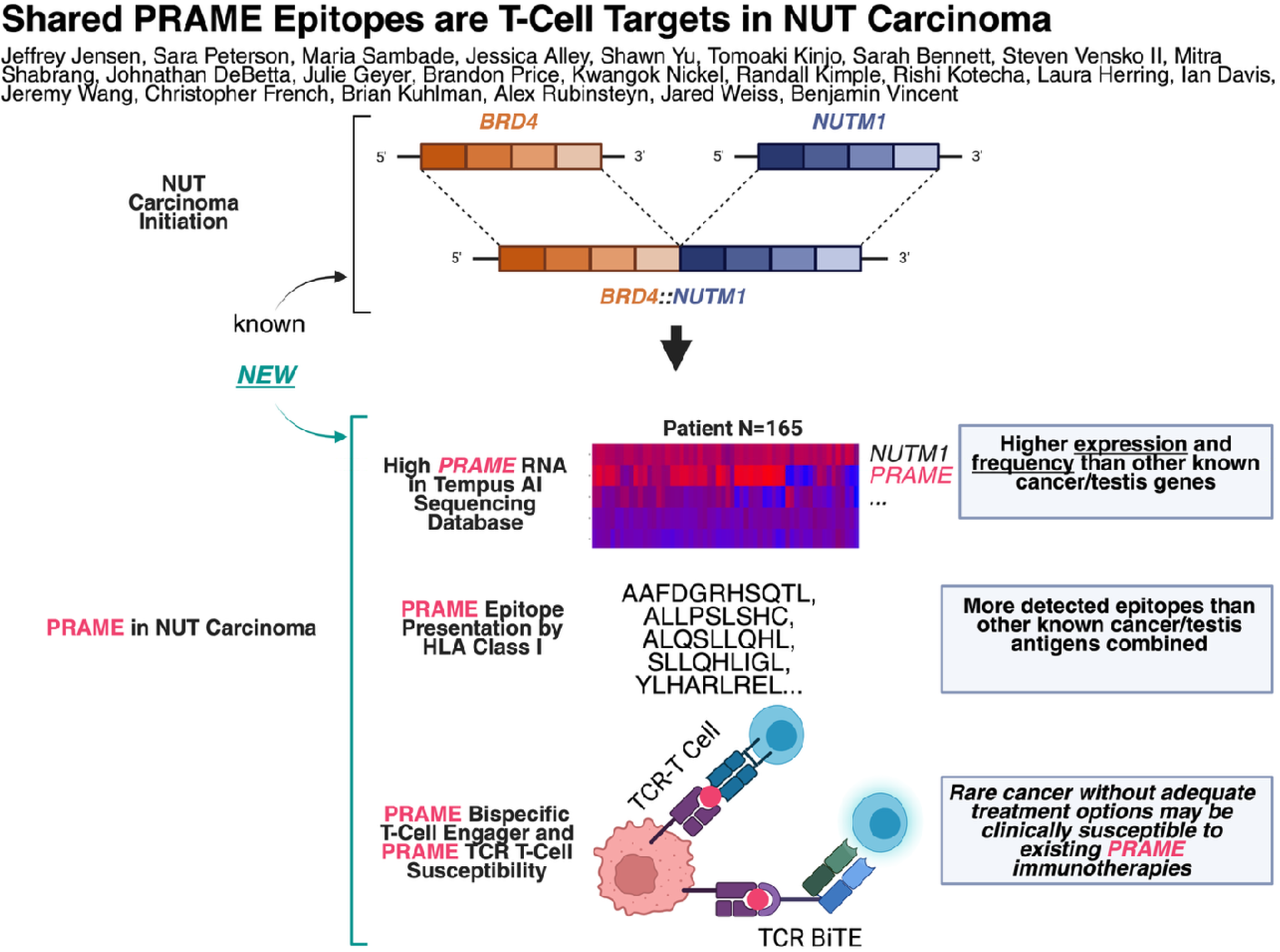

## Background

NUT carcinoma is a rare but underdiagnosed and devastating malignancy defined by ectopic expression of the Nuclear Protein in Testis gene (“*NUT*” or “*NUTM1*”), most commonly through an in-frame translocation to the bromodomain-containing *BRD4* with formation of the *BRD4::NUTM1* fusion gene (75% of cases) (1). BRD4::NUTM1 is a potent oncoprotein that initiates NUT carcinoma by binding to acetylated H3K27 (“H3K27ac”), recruiting p300 and catalyzing the hyperacetylation of adjacent histones resulting in extended hyperacetylated domains (“megadomains”) with the corresponding overexpression of corresponding oncogenes, including *MYC* and *SOX2* (2, 3). NUT carcinoma was formerly labeled as NUT Midline Carcinoma owing to a predilection for afflicting midline structures and is now known to occur most commonly in the lungs and sinonasal passages (4). NUT carcinoma is not associated with smoking or any known exposures and afflicts individuals of all ages including infants, children, and young adults (5-7). Clinical outcomes in adults are dismal with median survival after diagnosis of 6.7 months and similar outcomes in children due to rapid tumor growth, high metastatic potential, and lack of sensitivity to systemic therapies (4, 8). Treatment for localized disease is focused on aggressive attempts at local control with wide local resection and adjuvant radiation and/or chemotherapy. However, even among NUT carcinoma patients treated with complete surgical resection of the initial tumor, outcomes are poor. In one cohort, patients treated with surgical resection experienced a median survival time of 415 days post-diagnosis with no patient surviving for more than 3 years post diagnosis (4, 9).

Targeted therapeutic strategies to date have focused on blocking the binding of BRD4 to H3K27ac using bromodomain inhibitors, but in phase I/II clinical trials these inhibitors have yielded progression free survival times of only 1-2 months (10-13). At present, all clinical trials testing new therapeutics for NUT carcinoma patients in the United States are testing bromodomain inhibitors (NCT05019716, NCT05372640, and NCT05488548). NUT carcinoma tumors are PD-L1 low, tumor mutational burden low, and immune “cold” tumors that are also highly refractory to cytotoxic chemotherapy and T-cell checkpoint blockade (4). New treatment modalities are clearly needed.

Interest in treating challenging solid tumors with HLA-restricted TCR-based therapies is growing. The first phase III clinical trial of a TCR x CD3 activator bispecific (also known as an ImmTAC or BiTE) showed overall survival benefit in the frontline setting for HLA-A*02+ uveal melanoma targeting the gp100 peptide (14). Recently, the SPEARHEAD-1 trial showed durable responses in patients with heavily pre-treated synovial sarcoma or myxoid round cell liposarcoma with afamitresgene autoleucel, a TCR T-cell therapy expressing a TCR targeting an HLA-A*02 bound epitope from the cancer/testis gene *MAGE-A4* (15). This resulted in the first TCR T-cell product approval for afamitresgene autoleucel and was quickly followed up by the IGNYTE-ESO trial which similarly showed durable responses in the same disease types with letetresgene autoleucel, another TCR T-cell product targeting epitopes from the cancer/testis gene *NY-ESO-1* (HLA-A*02:01, -A*02:05, or -A*02:06) (16).

Like NUT carcinoma, synovial sarcoma and myxoid/round cell liposarcoma are uncommon fusion protein-driven malignancies. >90% of these sarcomas harbor the pathognomonic t(X;18) or t(12;16)(q13;p11), respectively (17, 18). Whereas MAGE-A4 and NY-ESO-1 antigens are common in these sarcomas, the antigenic landscape of NUT carcinoma and the susceptibility of NUT carcinoma to T-cell mediated therapeutics are unknown. We hypothesized that if BRD4::NUTM1 causes dysregulated gene transcription through reliable epigenetic changes, then BRD4::NUTM1 could also upregulate genes encoding cancer specific antigens that would serve as actionable TCR targets. Here we show that BRD4::NUTM1 causes transcription of the cancer/testis gene *PRAME*, and that *PRAME* is the predominant cancer/testis gene expressed within a clinical patient cohort of *NUTM1* fusion genes and especially in patients with canonical NUT carcinoma fusions (*BRD4::NUTM1*, *BRD3::NUTM1*, *NSD3::NUTM1*) (19). We also show that NUT carcinoma cells present PRAME epitopes with HLA Class I molecules and are susceptible to T-cell mediated killing mediated by an affinity-matured PRAME epitope TCR x CD3 activating bispecific molecule or by αPRAME TCR T-cells, with both modalities generated from TCRs currently in clinical testing for common cancers. Our work identifies PRAME as a promising target for cancer vaccines, TCR-based therapies, and TCR-mimetics in NUT carcinoma phase I/II clinical trials.

## Methods

### Reagents

Cell lines were maintained in DMEM with 10% FBS and 1% penicillin/streptomycin. The PDX donor signed consent version 12 of the “Consent Form for Use of Tissue, Blood, and Other Biological Materials in Research and Authorization for Use of Identifiable Health Information in Biobanking” for the University of Wisconsin Carbone Cancer Center Translational Science BioCore which was IRB approved on 10/21/2015. The patient’s tumor and a deidentified pathology report were provided to our investigators. The tumor was chopped and minced into small pieces, dissolved in media containing FBS (10%), penicillin/streptomycin (1%), amphotericin B (1%), and matrigel (1:1 mixture with tissue prep) and injected through a 21-gauge needle into the flanks of female NSG mice. The tumor was then propagated in the flanks of NSG mice (using the technique above) and purified for analyses using tumor dissociation and mouse cell depletion kits (Miltenyi Biotec). RNA was isolated using RNEasy columns (Qiagen). Western blots used PVDF and blocking in 5% milk. Antibodies are described in **Supplementary Table 1**. siRNA transfections were performed with Lipofectamine RNAiMAX and 10 μM pools of four siRNAs synthesized by IDT with [dT][dT] overhang, desalted purification, and no 3’ modifications (**Supplementary Table 2**). Tet-on, blasticidin selection lentiviral pInducer20 vectors containing *BRD4::NUTM1* or *GFP* have been described (20). VSV-G pseudotyped lentiviral particles were packaged in 293T cells and purified by filtration. TMA immunohistochemistry was done by BWH Dana-Farber-Harvard Medical School Specialized Histopathology Services-Longwood (SHL) Core using αPRAME (Biocare Medical ACI3252B, 1:100).

### Immunopeptidome Predictions

Antigen prediction used LENS v1.5.1 (21). Sample-level FASTQs were trimmed and filtered for quality using fastp 0.23.1 (22). The resulting FASTQ pairs were aligned to a modified hg38 reference genome (excluding the Epstein-Barr Virus (EBV) contig) using STAR 2.7.0f with parameters *--quantMode TranscriptomeSAM --outSAMtype BAM SortedByCoordinate --twopassMode Basic –outSAMunmappedWithin (23)*. Transcript-level quantifications were generated using Salmon 1.1.0 with a GENCODE v37 GTF, augmented with human retroviral-like sequences from gEVE (24). CTAs were defined using a manually curated list of genes with testis and placenta-specific expression (**Supplementary Table 3**). The initial list of CTAs was pulled from CTDatabase and manually filtered using Human Protein Atlas.

Splice variants were detected using SNAF 0.7.0 (25). Expressed viruses were detecting using the VirDetect workflow. HLA typing for class I alleles was performed using seq2HLA 2.2 (26). We processed protein sequences from CTAs in the 50th expression percentile for downstream peptide prediction. Gene fusions were predicted using STARfusion 1.10.1 with recommended parameters and trimmed RNA FASTQs. Relevant protein sequences derived from CTAs, fusion, splice variants, viruses, and ERVs were filtered for potential peptides of interest using NetMHCpan 4.1b (27).

### RNA Sequencing

Library preparation used Illumina’s Stranded mRNA Prep Ligation kit, starting with 125 ng of RNA. The average fragment length for these libraries was between 300-400 bp. Paired-end sequencing was performed using the NovaSeq 6000 version 1.5 (2X100), obtaining at least 45M reads per sample. Bulk RNA-seq samples were processed using selected components of the nf-core RNA-seq pipeline (v3.14), an open-source workflow managed with Nextflow. Reads were demultiplexed, and quality was assessed using FastQC, with results summarized in MultiQC (28). Adapter trimming was performed with cutadapt, ribosomal RNA was removed with SortMeRNA, followed by alignment to the GRCh38 reference genome using STAR (29). Transcript-level quantification was conducted with Salmon (24). Counts were summarized to the gene level and normalized to TPM to generate a gene x sample matrix. All downstream visualization and analysis were performed in R. All scripts used for this project are publicly available on GitHub (https://github.com/pirl-unc/PRAME_in_NUT_Carcinoma).

### Immunopeptidomics and Targeted Mass Spectrometry

Snap frozen cell pellets were submitted to Biognosys for immunopeptidomics. Samples and a processing quality control sample (JY cells) were lysed using Biognosys’ Lysis Buffer for 1 hour at 4°C. After removal of cell debris, the lysates were incubated with bead coupled antibodies (clone W6/32) to capture HLA class I complexes with associated peptides. Bound protein complexes were washed and peptides eluted using Biognosys’ optimized immunopeptidomics protocol. Clean up for mass spectrometry was carried out using an Oasis HLB µElution Plate (30 µm). Immunopeptides were dried down using a SpeedVac system and resuspended in 10 µl of 1% acetonitrile and 0.1% formic acid in water. Samples were pooled and fractionated into 5 fractions using high pH reversed phase fractionation (HPRP) for spectral library generation.

For DDA LC-MS/MS measurements of fractionated samples, 8 µl of immunopeptides per sample were injected to a Biognosys in-house packed reversed-phase column on a NeoVanquish UHPLC nano liquid chromatography system connected to a Orbitrap Exploris 480 mass spectrometer equipped with a Nanospray Flex ion source and a FAIMS Pro ion mobility device. LC solvents were A: water with 0.1% FA, B: 80% acetonitrile. The nonlinear LC gradient was 1 - 40% solvent B in 120 minutes followed by a column washing step in 90% B for 11 minutes, and a final equilibration step of 1% B for 5 minutes at 64°C with a flow rate set to 250 nl /min. MS1 precursor scans were acquired between 330-1250 m/z at 120,000 resolution with data dependent MS2 scans at 60,000 resolution. MS2 scans were acquired in cycle time of 1.6 s and 1.4 s per applied FAIMS compensation voltage. Only precursors with charge state 1-5 were isolated for dependent MS2 scans.

For DIA LC-MS/MS measurements of study samples, 8 µl of immunopeptides per sample were injected to a Biognosys in house packed reversed phase column on a NeoVanquish UHPLC nano liquid chromatography system connected to a Orbitrap Exploris 480 mass spectrometer equipped with a Nanospray Flex ion source and a FAIMS Pro ion mobility device (Thermo Scientific). The system was used in trap and elute configuration using a 300 µm x 5 mm PepMap Neo C 18 Trap 3 PK cartridge pre column (Thermo Scientific 174502). LC solvents were A: water with 0.1% FA, B: 80% acetonitrile, 0.1% FA in water. The nonlinear LC gradient was 1-40% solvent B in 120 minutes followed by a column washing step in 90% B for 11 minutes, and a final equilibration step of 1% B for 5 minutes at 64° C with a flow rate set to 250 nl /min. The FAIMS DIA method consisted of one full range MS1 scan and 20 DIA segments as adopted from Bruderer et al (30) and Tognetti et al (31).

The shotgun and HRM mass spectrometric data was analyzed using Spectronaut v18, the false discovery rate on peptide and protein level was set to 1 %. A human UniProt .fasta database (Homo sapiens, 2025-01-01) appended with *NUTM1* fusion proteins was used for the search engine, also supplemented with mutated *NUTM1* and *PRAME* isoform fastas, allowing for non-specifically digested peptides (no trypsin cleaved termini) with a length between 7 and 12 amino acids, max. 2 variable modifications (N-term acetylation, carbamidomethylation [C], pSTY) and requiring carbamidomethylation methionine oxidation as fixed modification. The identification results were then exported as a hybrid spectral library. Raw HRM mass spectrometric data were analyzed using Spectronaut v18 with Qvalue filtering and no background signal integration or imputation. The hybrid spectral library generated in this study was used. Peptide level false discovery rate control was 1 % and cross-run normalization using global normalization to the median. MS2 spectra were visualized and annotated using the Interactive Peptide Spectral Annotator. Fragment ion tolerance was set to 5ppm for a-, b-, and y-ions.

For targeted mass spectrometry, cells were homogenized using a Precellys homogenization device (Bertin Technologies) and then lysed as above along with a quality control sample pellet of 10 million Raji cells. Protein concentrations were measured with a BCA assay (Pierce, Thermo Scientific) then HLA Class I-bound peptides were isolated, cleaned up, dried down and resuspended as above. Four stable isotope labeled reference peptides were spiked into the final peptide samples at known concentrations (manufactured by SciTide with quality grade +/-10% quantification precision, >95% purity). For LC-PRM measurements, 50% of total immunopeptide yield per sample (1 μg total peptide anticipated per injection) was injected with 5 fmol SIS peptides on custom-length Aurora 3 Freontier-type reversed phase column (IonOpticks) on a ThermoScientific NeoVanquishUHPLC nano-liquid chromatography system connected to a ThermoScientific Orbitrap Exploris 480 mass spectrometer equipped with a Nanospray Flex ion source and a FAIMS Pro ion mobility device (ThermoScientific). LC solvents were A: water with 0.1% FA; 80% acetonitrile, 0.1% FA in water. The LC gradient was 0-50% solvent B in 55.8 min followed by 51-90% B in 10 sec, 90% B for 5.9 min to wash the column and a 1.2 min step at 1% B to equilibrate the column. The total gradient length was 63.3 min. An unscheduled run in PRM mode was performed before data acquisition for retention time calibration as previously described (32). Signal processing and data analysis was carried out using SpectroDive 12.4 and Q-value filter of 1%.

### Bispecific Protein Production

Coding sequences for PRAME soluble TCR specific for the HLA-A*02:01-restricted SLLQHLIGL epitope were derived from sequences previously described for the PRAME Vα and Vβ domains and from the Protein Data Bank ID 6U07 crystal structure for the computationally thermostabilized Cα and Cβ domains (33, 34). Chimeric SP-34 αCD3 heavy and light chain sequences were published previously (35). One primary fusion construct and two auxiliary constructs were generated as clonal genes in mammalian expression vector plasmids (Twist Bioscience). The primary fusion constructs consisted of the SP-34 αCD3 heavy chain, a (Gly_4_Ser)_4_ linker, PRAME TCR Vα/Cα, and an 8xHis-Tag. The first auxiliary coding sequence encoded for SP-34 αCD3 light chain; the second for PRAME TCR Vβ/Cβ (**Supplementary Table 4)**.

All three plasmid constructs were co-transfected for transient expression in Expi293F cells cultured with Expi293 Expression Medium at 50 mL scale using ExpiFectamine 293 Transfection Kit, similarly to previously described with the following modifications (36). Plasmids for primary fusion construct, first auxiliary construct, and second auxiliary construct were mixed in a 3:1:1 mass ratio, totaling 1 µg plasmid mixture per milliliter of cell culture. At 18 hours after transfection, enhancer 1 (300 µL at 50 mL scale) and enhancer 2 (3 mL at 50 mL scale) and 6 mL of 3.5% (w/v) Hyclone Cell Boost 1 were added to the cells. Cell culture supernatant was harvested at 96 hours after transfection by pelleting cells at 3428 RCF at 4°C for 15 min and further clarified using a 0.22 µm polyvinylidene difluoride membrane filter (Millipore). Assembled PRAME bispecific constructs were affinity purified by batch-binding clarified medium with pre-equilibrated penta-Ni resin (Marvelgent) for 1 hour at 4°C and then transferred into a column to collect the resin. A three-step wash was performed by gravity at room temperature (RT) to eliminate protein contaminants. The first wash was performed with 6 CV (column volume) of high-salt Tris buffer [50 mM Tris (pH 8.0) and 1 M sodium chloride with 10 mM imidazole]; the second was performed with 2 CV wash with 1× PBS (pH 7.4) with 20 mM imidazole; and the third was performed with 2 CV wash with 1× PBS (pH 7.4) with 40 mM imidazole. Bound proteins were eluted in one step using 5 CV of 1× PBS (pH 7.4) and 500 mM imidazole to collect the protein and then sequential concentration at 4°C using a 30K MWCO Amicon-4 followed by an Amicon-0.5 centrifugal filter at maximum allowed RCF for each filter, serially exchanged into 1× PBS (pH 7.4) to ensure at least 1:1000 dilution of imidazole. Purity was assessed by SDS-polyacrylamide gel electrophoresis.

### TCR T-Cells

TCR T-cells were generated using a previously described approach to knock-out endogenous α and β chains, followed by retroviral transduction to express the anzutresgene autoleucel αPRAME TCR (37-39). CD8+ T-cells were isolated from healthy donor PBMCs using an EasySep Human CD8+ T-Cell Isolation Kit (StemCell Technologies) and stimulated for 72 hours with plate-bound αCD3 (1 μg/mL, BioLegend) and αCD28 (1 μg/mL, BioLegend) with Il-2 (300 IU/mL, StemCell Technologies), IL-7 (5 ng/mL, StemCell Technologies), and IL-15 (5 ng/mL, StemCell Technologies). Following activation, T-cells were electroporated with TCR α and β chain-targeting ribonucleoproteins using a 4D Nucleofector X (Lonza) and cultured for 48-72 hours with 180 IU/mL IL-2. To generate retroviral supernatants containing the TCR of interest, HEK292T cells were transfected with retroviral plasmids, including SPR plasmid containing the αPRAME TCR, using GeneJuice Transfection Reagent (Sigma-Aldrich) and harvested at 48 and 72 hours as previously described (39). TCR null CD8+ T-cells were transduced in Retro-Nectin-coated plates (Takara Bio) according to the manufacturer’s protocol and cultured for a minimum of 72 hours prior to screening. To confirm TCR expression and specificity, TCR T-cells were stained on ice with peptide-loaded tetramers (Flex-T A02 monomers, PE, BioLegend), αCD8 (BV421, Clone: Hit8a, BioLegend) and fixable live/dead stain (NIR, ThermoFisher). Cells were fixed and run on a BD Symphony A3 flow cytometer (BD Biosciences); analysis was performed with FlowJo (v.10.10.0).

### Cytotoxicity Assays

Whole blood was obtained from Gulf Coast Regional Blood Center. PBMCs were isolated using Ammonium-Chloride-Potassium Lysing Buffer and Density Gradient Medium (Stemcell Technologies). T-cells were isolated using a negative selection, magnetic bead kit (Stemcell Technologies) and then incubated with RPMI and 10% FBS for 5 days in an αCD3 coated (5 μg/mL, BD Biosciences) flask with carrier-free IL-2 (2 ng/mL, 18.2 IU/mL, R&D Systems) and αCD28 (2.5 μg/mL, BD Biosciences). Expanded T-cells were co-cultured in RPMI media with 10% FBS. Luminescence output was either read using CellTiter-Glo reagent for luciferase-free cell lines or by adding luciferin (250 μg/mL, Promega). Peptide pulses used 1 μg/mL of SLLQHLIGL peptide, synthesized by the UNC High-Throughput Peptide Synthesis and Array Facility over three hours (T2 peptide pulse) or 10 μg/mL of SLLQHLIGL (GenScript) added at the beginning of the experiment (TCR T-cell cytotoxicity).

## Results

### RNA sequencing and computational predictions suggest PRAME and NUTM1 are sources of HLA Class I peptide ligands in NUT carcinoma

To test the hypothesis that transcriptional dysregulation due to NUTM1 fusion oncoproteins would lead to the production of cancer-specific antigens, we obtained five NUT carcinoma cell lines (HLA-A*02 positive:TC-797, 10-15, 14169, PER403. HLA-A*02 negative: JCM1) and one HLA-A*02+ PDX described in **Table 1**. We then performed RNA sequencing on these samples and applied the LENS computational workflow (**Figure 1A**). LENS is an open-source method that predicts tumor-specific and tumor-associated antigens from genomic data (21). LENS predicted HLA Class I binding for peptides derived from several CTAs but only PRAME and NUTM1 had predicted epitopes across multiple samples (**Figure 1B-C**). Interestingly, LENS predicted the translationally relevant SLLQHLIGL (PRAME_425_) epitope in every sample except for TC-797 in which *PRAME* was transcriptionally silent (40, 41). Whole genome long-read DNA sequencing failed to identify a genetic cause for the absence of PRAME expression in TC-797 (data not shown). In screening cancer/testis genes using our manually curated list of CTAs (**Supplementary Methods Table 3**), out of 164 different cancer/testis genes only *PRAME* and *NUTM1* were consistently expressed across the tested samples (**Figure 1D**). To evaluate protein-level heterogeneity of PRAME we stained the samples with αPRAME and confirmed TC-797 to be PRAME- with PER403, 14169, and JCM1 showing high proportions of PRAME+ cells (**Figure 1E**). We concluded that in this N=6 set of NUT carcinomas, PRAME and NUTM1 are the most promising sources of cancer-specific antigens for therapeutic action.

### RNA sequencing and immunohistochemistry identify high frequency and levels of PRAME RNA and PRAME protein in NUT carcinoma patient samples

To determine if the predominance of *PRAME* and *NUTM1* expression among cancer/testis genes was reflected in patient samples, we searched the de-identified Tempus multimodal database for patients with detected *NUTM1* fusion genes (identified via Tempus xT [solid-tumor DNA] (42), Tempus xR [solid-tumor RNA] (43), or Tempus xF [circulating tumor DNA] assays) (44). Research filters below clinical reporting were applied to obtain patients with evidence of a *NUTM1* fusion, defined as containing at least 10 total supporting reads from DNA or at least 5 total supporting reads from RNA. Subject to these thresholds, 165 patients had evidence of *NUTM1* fusions, with a total of 33 distinct fusion partners (**Table 2**). Consistent with their role as canonical NUT carcinoma fusion partners, *BRD4::NUTM1, NSD3::NUTM1,* and *BRD3::NUTM1* accounted for the majority of cases (35.8%, 17.6%, and 10.9% respectively) with the remaining samples harboring one of the 30 other fusion partners (**Supplementary Figure 1A).** As expected, *BRD4::NUTM1*, *BRD3::NUTM1*, and *NSD3::NUTM1* fusions primarily occurred in the lung and head/neck primary sites – typical anatomic locations for NUT carcinoma (9). Analysis of gene expression from 162 of the patients with available RNA data revealed that out of 157 CTA genes covered in the Tempus xR assay, *PRAME* was identified as having the highest average expression after *NUTM1* (**Figure 2A**). We observed a bimodal distribution of *PRAME* expression in comparison to *NUTM1* across the cohort, designating *PRAME*^high^ and *PRAME*^low^ groups using a threshold of 3.1 log_2_(TPM+1) (**Figure 2B**). We found the *BRD4::NUTM1* group to be significantly enriched in *PRAME*^high^ samples (fisher exact, *p*<0.001) with *BRD3::NUTM1* and *NSD3::NUTM1* also having more *PRAME*^high^ samples than *PRAME*^low^ (**Figure 2C**). Of the remaining non-canonical fusion partners, 14 partners had at least one instance of *PRAME*^high^ with 3 fusion partners having more than one: *MGA::NUTM1*, *SLC12A6::NUTM1*, and *CIC::NUTM1* (**Supplementary Figure 1B**). Notably, only 1 of 13 *YAP1::NUTM1* samples was classified as *PRAME*^high^. Using the *PRAME*^high^ and *PRAME*^low^ groups, we identified several primary sites with the highest percentage of *PRAME*^high^ samples (**Figure 2D**). These frequently *PRAME*^high^ sites included typical NUT carcinoma sites of origin including sinuses, head/face/neck, the nasal cavity, and lung.

To assess for intratumoral heterogeneity of protein level PRAME expression in NUT carcinoma patient samples, we stained a previously described NUT carcinoma tissue microarray for PRAME and quantified the percentage of PRAME positive cells within the tumor (**Figure 2E, 2F**) (45). We found high frequency of PRAME expression in 28/77 (36%) of patient samples with ≥75% of tumor cells staining positive. Overall 43/77 (56%) of patient samples exhibited some PRAME staining.

### BRD4::NUTM1 drives PRAME expression

Due to the high levels and frequency of *PRAME* observed in cell lines (N=5), a PDX (N=1), the Tempus AI sequencing database (N=165), and the NUT carcinoma tissue microarray (N=77), and because PRAME expression is regulated by epigenetic changes including H3K27 acetylation in non-malignant human physiology (46), we hypothesized that *NUTM1* fusion oncoproteins might be contributing to transcriptional dysregulation and causing the ectopic expression of PRAME. To test this hypothesis, we transduced HEK 293T cells with tet-on *BRD4::NUTM1* or *eGFP* lentiviral particles and observed an increase in PRAME protein levels with *BRD4::NUTM1* expression but not with GFP (**Figure 3A**). We then used siRNAs targeting *NUTM1* to knockdown BRD4::NUTM1 in the five NUT carcinoma cell lines and immunoblotted for PRAME (**Figure 3B**). We found that in all cell lines except for the *PRAME* negative TC-797 cell line, BRD4::NUTM1 loss was accompanied by a decrease in PRAME protein levels. The decrease in PRAME with BRD4::NUTM1 knockdown was marginal in 14169 compared to other NUT carcinoma cell lines, so we performed a western blot time course in 14169 cells which revealed a greater effect on PRAME levels at a later timepoint, 4 days instead of 3 days, after *NUTM1* siRNA transfection (**Figure 3C**). We considered whether these effects might be mediated through the corresponding loss of MYC after BRD4::NUTM1 knockout, but in our review of the literature describing MYC-target genes we could find no evidence to suggest that *PRAME* is a MYC-target gene (47). We concluded that BRD4::NUTM1 is contributing to the expression of PRAME in *BRD4::NUTM1* NUT carcinomas, which may be independent of MYC.

### Immunopeptidomics on NUT carcinoma samples identifies more HLA Class I PRAME ligands than all other analyzed CTAs combined

We hypothesized that HLA Class I-bound PRAME epitopes would be detectable using immunopeptidome profiling with mass spectrometry. Shotgun (DIA) mass spectrometry was performed with the five cell lines and with the PDX to identify and characterize NUTM1, PRAME, and other CTA epitopes being presented by HLA Class I molecules and to guide therapeutic TCR selection. We identified a total of 33,644 unique peptides among the six samples **(Supplementary Figure 2**). Focusing on the presented CTAs, we identified 14 unique peptides from 6 proteins out of all CTAs with detection in at least 1 sample. Of these 14 peptides, 9 corresponded to PRAME epitopes and 1 corresponded to a novel, previously undescribed NUTM1 epitope (**Figure 4A**). The remaining four CTA peptides corresponded to *MAGE-A1*, *CT83*, *CTAG2* and *ACTL8*. As expected, no PRAME epitopes were discovered in the *PRAME* negative TC-797 cell line. The well described and therapeutically actionable HLA-A*02:01/SLLQHLIGL (PRAME_425_) epitope was detected in the HLA-A*02+ PER403 and PDX samples but not 10-15 or 14169 (48-50). To detect fusion junctional peptides, long read RNA sequencing was performed and used to generate consensus fusion sequences (**Supplementary Figure 3**). A reference human proteome augmented with these fusion sequences failed to identify junctional fusion peptides.

Due to differential fragmentation of amino acids, quantification of peptide epitopes with untargeted immunopeptidomics is inherently semi-quantitative. Because SLLQHLIGL (PRAME_425_) was only detected in 2/4 (50%) of PRAME+, HLA-A*02+ samples and because the SLLQHLIGL (PRAME_425_) epitope with the most clinical developmental interest had lower semi-quantitative signal intensities than YLHARLREL (PRAME_462_), we developed targeted mass spectrometry assays for three PRAME epitopes and one novel NUTM1 epitope using single- and double-labeled peptide standards to quantify these epitopes across our six NUT carcinoma samples. Single- or double-labeled standards were used to enhance our confidence in the quantitative results across two independent sets of experiments. The labeled peptide standards used for these assays are described in **Table 3**. We found similar results and trends using single- and double-labeled peptide standards, and were able to detect SLLQHLIGL (PRAME_425_) in 4/4 (100%) of PRAME+, A02+ samples at abundances higher than PRAME_312_ or PRAME_462_. Interestingly, DVYENFRQW (NUTM1_237_) (**Figure 4B-D**) exhibited the highest absolute abundance of all epitopes and samples in the JCM1 cell line in the single-labeled assays and the second highest absolute abundance in the JCM1 cell line in the double-labeled assays but was very low abundance or non-detectable in the other 5/6 samples, likely due to DVYENFRQW being a HLA-A*25:01 ligand (MHCflurry predicts this as the only JCM1 HLA class I molecule with binding specificity for DVYENFRQW).

Based on the increased prevalence of PRAME epitopes in our samples relative to NUTM1, the rarity of HLA-A*25:01 compared to HLA-A*02:01, and based on the availability of existing PRAME therapeutics for testing, we focused on therapeutic targeting of PRAME epitopes in NUT carcinoma.

### PRAME+ NUT carcinoma cells are susceptible to T-cell killing mediated by an αPRAME_425_ x SP34 αCD3 BiTE or by αPRAME_425_ TCR T-cells

The susceptibility of NUT carcinoma to TCR-mediated therapeutics like TCR bispecifics or TCR T-cell therapies is unknown. To explore this issue and to evaluate the translational relevance of PRAME cancer-specific antigens, we created an αPRAME_425_ TCR x SP34 αCD3 BiTE molecule with the same variable regions as the affinity-matured TCR BiTE brenetafusp (**Figure 5A**). Brenetafusp has completed phase I/II testing and is currently being evaluated as first line therapy in a randomized phase 3 trial (NCT06112314) for melanoma patients (51). Because our αPRAME_425_ TCR x SP34 αCD3 BiTE has significant homology to brenetafusp including perfect homology to the variable regions of the TCR, we anticipated our bispecific molecule to have specificity for the same SLLQHLIGL (PRAME_425_) epitope brenetafusp targets. We used SLLQHLIGL peptide pulsing and TAP deficient T2 cells in co-culture with allogeneic T-cells to demonstrate enhancement of bispecific-mediated cytotoxicity with peptide pulse.

TAP deficient T2 cells have empty HLA molecules on the cell surface and are loaded by incubating the cells with an HLA peptide ligand. Killing of peptide-pulsed T2 cells thus demonstrated the specificity of the bispecific molecule for the SLLQHLIGL epitope (**Figure 5B**). We then titrated the concentration of the bispecific in the co-cultures of a *PRAME*+, HLA A02+ NSCLC cell line in the presence of allogeneic T-cells and found a dose-dependent effect (**Figure 5C**). The *PRAME* negative TC-797 and the *PRAME* positive 10-15 and 14169 cell lines were tested alongside a haplo-incompatible NSCLC cell line (H460, HLA A*24:02, *68:01) for T-cell mediated cytotoxicity in co-culture experiments using 10^-8^ M of bispecific molecule. We observed no cytotoxic effect of the bispecific in TC-797 or H460 at either of the tested effector:target ratios but we did observe potent cytotoxicity that was enhanced by a higher effector:target ratio in both *PRAME* positive NUT carcinoma cell lines (**Supplementary Figure 4A-B**).

This first cytotoxicity experiment was done using luminescent cell lines, however, PER403 and JCM1 luminescent cell lines were unavailable. Thus, we designed a second experiment to include PER403 and JCM1 in which T-cells were washed off of the adherent tumor cells and then tumor cell viability was assessed using CellTiter-Glo. To further demonstrate specificity of the αPRAME_425_ x SP34 αCD3 bispecific for HLA Class I-mediated cytotoxicity we also created a control, αHTLV-1 Tax x SP34 αCD3 bispecific molecule identical to the PRAME bispecific except for changes in the variable regions (**Figure 5A**). Again, we observed potent T-cell mediated cytotoxicity in PRAME+, A02+ NUT carcinoma cells with the αPRAME_425_ bispecific molecule but not with the αHTLV-1 Tax bispecific or in no bispecific controls (**Figure 5E, Supplementary Figure 4C**).

Lastly, to demonstrate that NUT carcinoma is likely susceptible to PRAME TCR immunotherapies as a class we also generated off-the-shelf αPRAME_425_ TCR T-cells using the TCR from anzutresgene autoleucel: a PRAME TCR T-cell product in early phase clinical testing for patients with melanoma and other advanced solid tumors (NCT03686124, NCT06743126). The native TCR was genetically ablated using transfection with Cas9 protein and TCR sgRNA guides (1 α chain sgRNA, 1 β chain sgRNA), and then TCR null T-cells were transduced with αPRAME_425_ TCR encoding lentiviral particles or with vehicle control (**Figure 5A**). Flow cytometry with αTCR immunoglobulin and SLLQHLIGL tetramer staining demonstrated successful ablation of the naïve TCR and lentiviral TCR transduction efficiencies of 66% or 41%, respectively by staining method (**Figure 5D, Supplementary Figure 5**). We then co-cultured TCR null T-cells and αPRAME_425_ TCR T-cells with the five NUT carcinoma cell lines, with and without SLLQHLIGL peptide pulse and found complete or nearly complete ablation of target tumor cells at the lowest E:T ratio tested (1:1) in all 3 PRAME+, HLA-A*02+ NUT carcinoma cell lines treated with αPRAME_425_ TCR T-cells but not with αHTLV-1 Tax TCR T-cells (**Figure 5F, Supplementary Figure 4D**). Importantly, in HLA-A*02:01 but PRAME- TC-797, peptide pulse with SLLQHLIGL peptide rescued the cytotoxicity effects of αPRAME_425_ TCR T-cells consistent with antigen-directed killing. We concluded that PRAME epitopes are a previously unrecognized therapeutic vulnerability in NUT carcinoma.

## Discussion/Conclusion

Recent advances in antigen-directed immunotherapy have resulted in a new generation of therapeutics that leverage cell surface protein or TCR/peptide/HLA binding interactions for tumor targeting and specificity. Approved agents in solid tumors target either HLA-presented peptide epitopes such as gp100 in uveal melanoma, MAGE-A4/NY-ESO-1 epitopes in synovial sarcoma/myxoid round cell liposarcoma, or surface targets such as DLL3 in small cell lung cancer (14, 15). To identify promising targeted immunotherapies for NUT carcinoma, a highly lethal and treatment-refractory malignancy, we performed RNA sequencing on a selection of NUT carcinoma samples and applied the results to the antigen prediction software LENS which predicted PRAME as a plentiful source of cancer-specific antigens in NUT carcinoma (21). We found that *PRAME* was highly expressed in cell lines and the PDX model, consistent with observations in a large cohort of *NUTM1* fusion patient samples (especially among canonical *NUTM1* fusion patient samples) as well as in a NUT carcinoma tissue microarray.

Using genetic approaches, we showed that elevated PRAME levels in *BRD4::NUTM1* NUT carcinoma are enhanced or at least maintained by BRD4::NUTM1. By applying immunopeptidomics to our panel of NUT carcinoma samples we also detected 9 PRAME epitopes, including the actionable SLLQHLIGL (PRAME_425_) epitope that is targeted by brenetafusp and anzutresgene autoleucel. NUTM1 epitopes were nearly absent from our immunopeptidomics results, likely due to immunoselection for non-antigenic fusion proteins, limitations of detection by shotgun immunopeptidomics, and/or strong nuclear localization of NUTM1 fusion proteins (13). However, we did identify a single novel NUTM1 epitope, likely binding HLA-A*25:01, confirmed through targeted mass spectrometry to be absent or at very low levels of abundance in all of the HLA-A*25:01 negative samples. Future identification of other NUTM1 epitopes for NUTM1-specific TCR therapeutic development will require more sensitive immunopeptidomic methods, or perhaps targeted mass spectrometry using computationally predicted epitopes.

PRAME epitopes are appealing candidates for peptide-HLA-directed therapeutics because of the strong localization of PRAME to the testis in healthy human physiology and because they arise as a direct result of the initiating genetic lesion in NUT carcinoma (*BRD4::NUTM1*). We created an αPRAME_425_ TCR x SP34 αCD3 molecule modeled after brenetafusp that then showed potent, T-cell-mediated cytotoxicity against *PRAME*+ NUT carcinoma cells but not *PRAME*- NUT carcinoma or haploincompatible NSCLC cells. We also showed NUT carcinoma cells to be susceptible to αPRAME_425_ TCR T-cell antigen-directed killing using the TCR from anzutresgene autoleucel, building our confidence that PRAME immunotherapies as a class might perform well against NUT carcinomas in clinical trials.

There are multiple limitations of this work. We demonstrated PRAME antigen-specific cytotoxicity *in vitro* have not presented *in vivo* treatment models here. This is in part because brenetafusp and anzutresgene autoleucel are already in clinical trials, making unexpected toxicities or dosing issues in mice less relevant. Secondly, mouse models for human antigen-directed therapeutics are contrived as they require human T-cell injections into immunocompromised mice and the human T-cells have alloreactivity to mouse tissues. We also encountered unexpected results in our cytotoxicity experiments. Specifically, the αPRAME bispecific enhanced proliferation in PRAME-TC-797 at E:T=5:1 in two independent experiments. The αHTLV-1 Tax bispecific also exhibited this effect. These data suggest that CD3 stimulation enhances TC-797 proliferation in the absence of antigen-binding, perhaps through cytokine-mediated effects. There was also unexpected cytotoxicity against HLA-A*02- JCM1 cells with both the αPRAME BiTE and αPRAME TCR T-cells even though SLLQHLIGL (PRAME_425_) is broadly recognized as an HLA A02 ligand. This may be due to reactivity against the SLLQHLIGL (PRAME_425_) peptide presented by HLA-B*08:01, as computational predictions suggest SLLQHLIGL (PRAME_425_) has high binding affinity for this JCM1 HLA-B allele (52).

In conclusion, we report here that the most common oncogene initiating NUT carcinoma, *BRD4::NUTM1*, enhances PRAME expression, that PRAME is a frequently and highly expressed cancer/testis gene in *BRD4::NUTM1, NSD3::NUTM1* and *BRD3::NUTM1* NUT carcinomas, and that the resulting PRAME epitopes are actionable targets for TCR therapeutics in NUT carcinoma. Our work adds NUT carcinoma to the growing cadre of malignancies for which there is immense interest in targeting PRAME. Specifically, PRAME is prevalent in genetically heterogeneous cancers including melanoma, acute myeloid leukemia, NSCLC, ovarian cancer, and others (53). In healthy human physiology, *PRAME* is turned on at the earliest stages of spermatogenesis and is silenced by histone modifications upstream of the translation initiation site (46). In malignancies, PRAME expression has been associated with worse clinical outcomes, DNA double strand breaks, aneuploidy, and genetic instability (46, 54-56). Oncogenic selection pressures in NUT carcinoma likely favor overriding the natural protections against ectopic PRAME production via BRD4::NUTM1-p300 acetylation of H3K27 (2, 45, 57). EZH2 is active in NUT carcinoma and promotes inhibitory trimethylation of H3K27 which is mutually exclusive with H3K27ac domains in NUT carcinoma (45). Thus, H3K27 trimethylation is a plausible mechanism by which TC-797 and other *BRD4::NUTM1* patient samples could restrict *PRAME* expression. Ultimately, further assays of histone states, such as ChIP-Seq or CUT&RUN, could explore this possibility. The mechanisms promoting transcription of *PRAME* are likely conserved between BRD4::NUTM1, BRD3::NUTM1, and NSD3::NUTM1 because BRD4 and BRD3 are highly homologous (>80%) and because NSD3::NUTM1 recruits BRD4, thus forming a protein complex with the same constituents as BRD4::NUTM1 (19, 58).

Our work suggests that clinical trials of PRAME targeting TCR therapeutics including TCR T-cell therapies, endogenous T-cell therapy, TCR BiTEs/ImmTACs, tumor vaccines, and/or immunoglobulins specific for HLA-bound PRAME epitopes (“TCR mimetics”) are warranted in NUT carcinoma. Potential agents to test include the aforementioned brenetafusp and the TCR T-cell therapy anzutresgene autoleucel (40, 41). A PRAME-specific TCR is included in the presently recruiting trial NCT05973487: a personalized TCR T-cell trial for patients with locally advanced or metastatic solid tumors. An RNA-loaded dendritic cell vaccine using *PRAME* has completed phase I testing for allogeneic stem cell transplant-ineligible acute myeloid leukemia and other PRAME vaccines are in development (59, 60). Pr20, a PRAME-specific TCR mimetic has been developed and showed activity against human leukemias in xenografts (61). Brenetafusp has been tested in PRAME+ and PRAME- tumors (as determined by IHC) with PRAME+ tumors showing better responses (40). Our clinical cohort showed common NUT carcinoma anatomical origins to be associated with very high or universal *PRAME* expression, suggesting that either IHC and/or anatomical origin could be reasonable inclusion criteria for PRAME immunotherapy trials in this population. Further, the growing set of PRAME therapeutics may have greater efficacy in NUT carcinoma than in melanoma, NSCLC or other common malignancies without unifying genetic etiologies because of a mechanistic link between the central genetic rearrangements in NUT carcinoma and *PRAME* expression. Mechanistic studies linking bromodomain fusion proteins to PRAME and clinical trials testing PRAME therapeutics in NUT carcinoma are logical next steps to discovering efficacious treatments for this rare, orphan disease.

## Supporting information

Tempus AI CTA Data

## Declarations

### Ethics approval and consent to participate

All genetic engineering experiments involving lentivirus or siRNAs were approved by the UNC Institutional Biosafety Committee. All animal experiments were approved by the UNC Institutional Animal Care and Use Committee. All patient samples analyzed in the Tempus AI Sequencing Database were deidentified, and patients consented for their deidentified data to be used in research.

### Consent for publication

All authors consent that contingent on the acceptance thereof, for this research to be published in JITC with all appropriate requirements of copyright and access as this journal demands.

### Availability of data and material

All scripts and non-genomic, non-mass spectrometry raw data used to generate the figures are available at GitHub (https://github.com/pirl-unc/PRAME_in_NUT_Carcinoma). RNA sequencing data from the five NUT cell lines and the patient derived xenograft will be available on Gene Expression Omnibus. A CSV file is available in supplementary data which includes gene expression data for cancer/testis genes in the Tempus AI patient cohort. All other sequencing data for this cohort is the property of Tempus AI and will be shared at their discretion. Any other raw data (including mass spectrometry, TMA PRAME staining images) not owned by Tempus AI are available upon request.

### Authors’ contributions

JLJ – Generated hypotheses, performed experimentation for all figures, wrote the first draft of the manuscript, revised the manuscript, and obtained funding

SKP – Generated hypotheses, performed sequencing analyses, produced the figures, wrote the first draft of the manuscript, and revised the manuscript

MaS – Produced western blot samples, produced western blots, and prepared samples for immunopeptidomics

JRA – Prepared samples for immunopeptidomics, performed bispecific and TCR T-cell cytotoxicity assays, and edited the manuscript

SY – Designed and produced αPRAME_425_ x SP34 αCD3 bispecific, designed and produced the αHTLV bispecific, performed testing in T2 cells and NSCLC cells, and revised the manuscript

TK - Designed and produced αPRAME_425_ x SP34 αCD3 bispecific, designed and produced the αHTLV bispecific, performed testing in T2 cells and NSCLC cells, and revised the manuscript

SNB – Designed and produced αPRAME TCR T-cells and performed flow cytometry staining to quantify transduction efficiency

SV – Predicted antigens using LENS, produced the corresponding figures, and also wrote the methods section for LENS

MiS – Analyzed flow cytometry data

JDD – Prepared samples for western blotting

JKG – Analyzed long read DNA sequencing

BAP – Curated the *NUTM1* fusion cohort in the Tempus AI Sequencing Database and performed sequencing analysis

KPN – Developed and provided the NUT carcinoma patient-derived xenograft

RJK - Developed and provided the NUT carcinoma patient-derived xenograft RSK – Developed and provided the NUT carcinoma cell line JCM1

LEH - Analyzed immunopeptidomics data, generated mass spectrum figures, and revised the mass spectrometry methods section.

IJD – Mentored SKP, assisted in sequencing analysis, and revised the manuscript

JRW – Mentored JKG and analyzed long read DNA sequencing

CAF – Provided NUT carcinoma cell lines, produced NUT carcinoma tissue microarray, stained stained the TMA for PRAME, and revised the manuscript

BK – Mentored SY and TK, designed and produced αPRAME_425_ x SP34 αCD3 bispecific, designed and produced the αHTLV bispecific, and revised the manuscript

JW – Conceived the project, mentored JLJ, generated hypotheses, revised the manuscript, and obtained funding

AR – Conceived the project, mentored SKP, generated hypotheses, designed experiments, analyzed sequencing data, revised the manuscript, and obtained funding

BGV – Conceived the project, supervised the project, mentored JLJ, generated hypotheses, designed experiments, revised the manuscript, and obtained funding

## Acknowledgments

We would like to acknowledge the work and contributions from multiple core facilities at UNC including the Immune Monitoring and Genomics Facility, the High Throughput Sequencing Facility, the Pathology Services Core Facility, the Peptide Synthesis Core Facility, and the lentiviral/shRNA core facility.

## List of Abbreviations

LENS: Landscape of Neoantigen Software
NSCLC: Non-small cell lung cancer
TCR: T-cell receptor
CTA: Cancer testis antigen
BiTE: Bispecific T-cell engager
ImmTAC: Immune mobilizing monoclonal T-cell receptors against cancer
DMEM: Dulbecco’s modified eagle medium
PDX: Patient-derived xenograft
TPM: Transcripts per million
RCF: Relative centrifugal force
IND: Investigational new drug
IHC: Immunohistochemistry
DDA LC-MS/MS: Data-dependent acquisition liquid chromatography tandem mass spectrometry
DIA: Data-independent acquisition
TMA: Tissue microarray

---

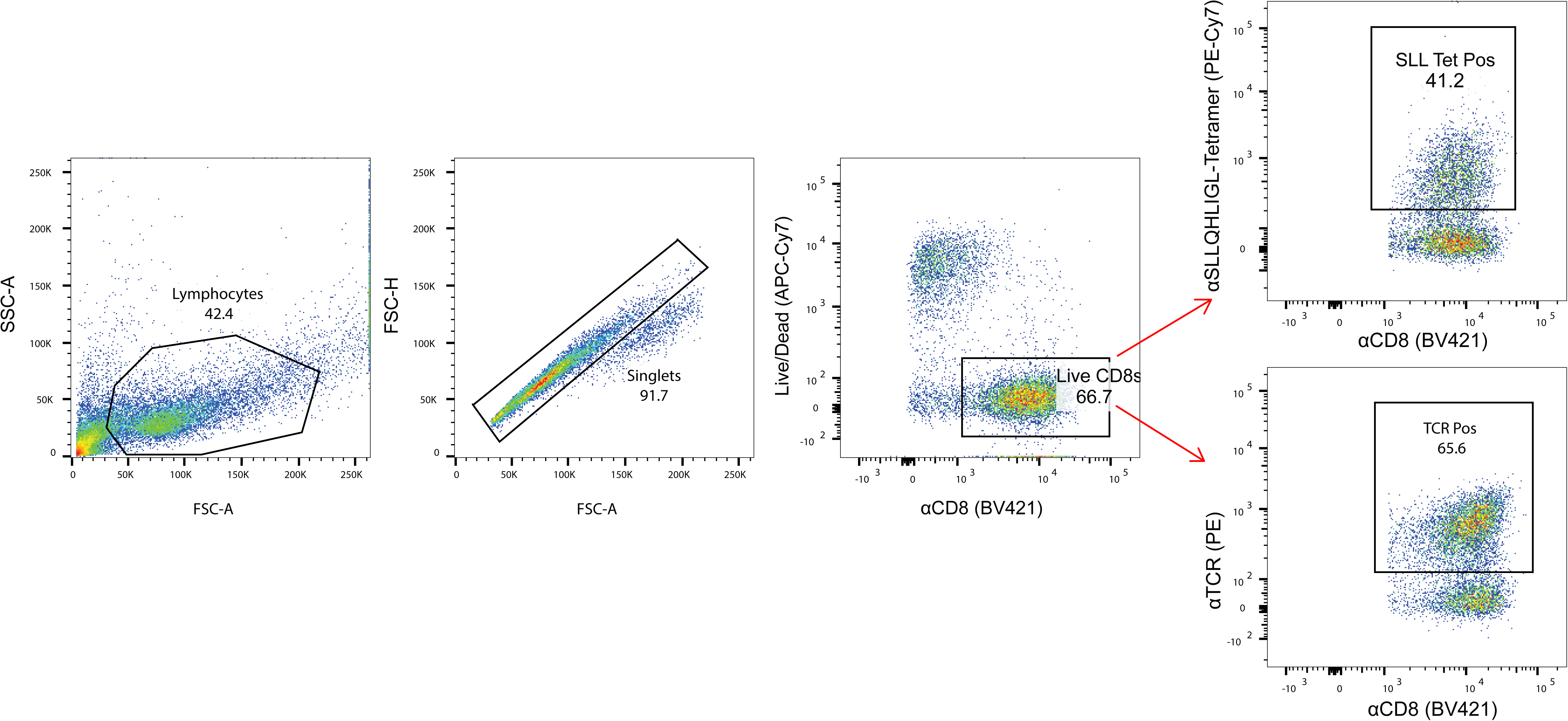

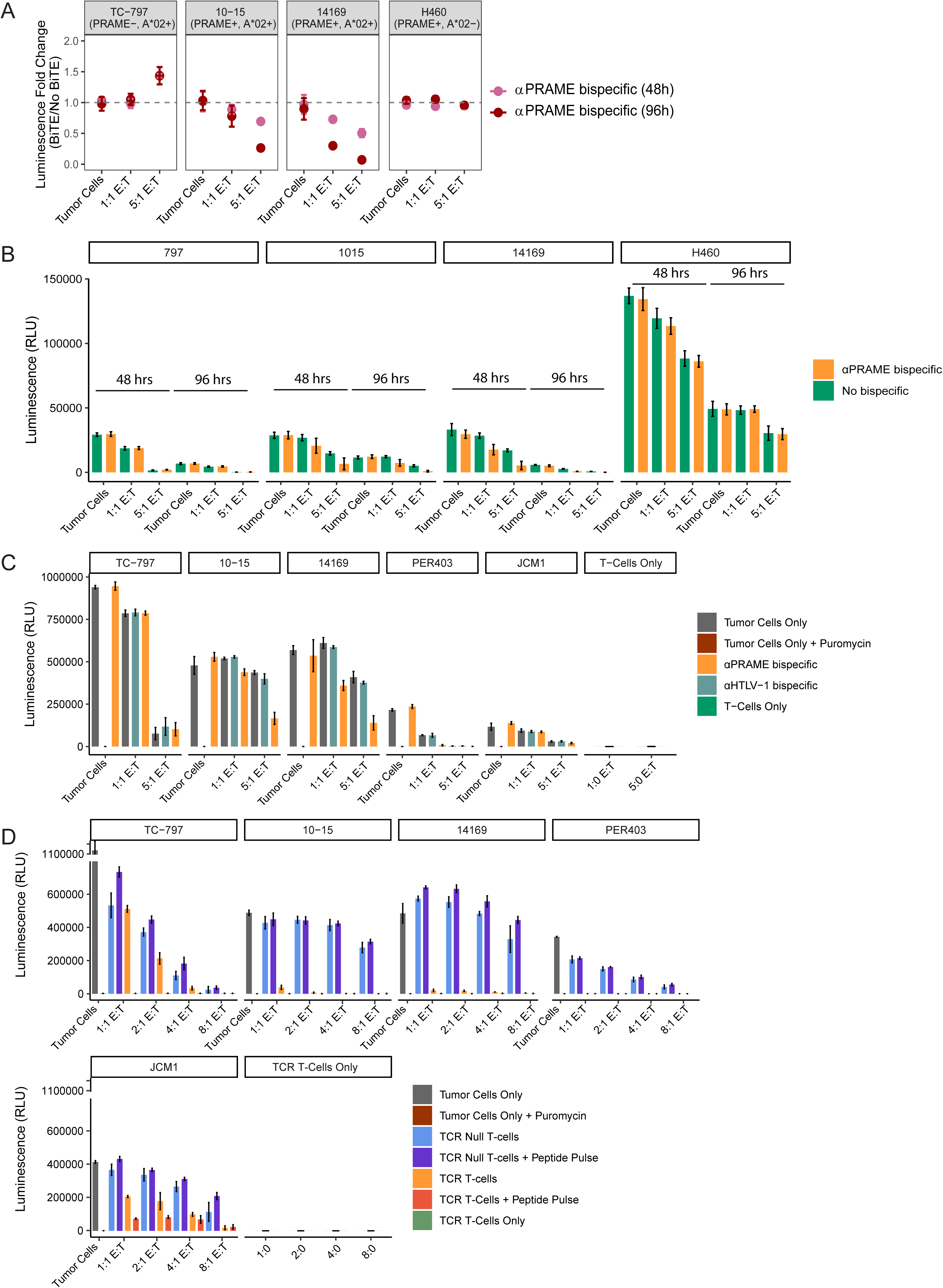

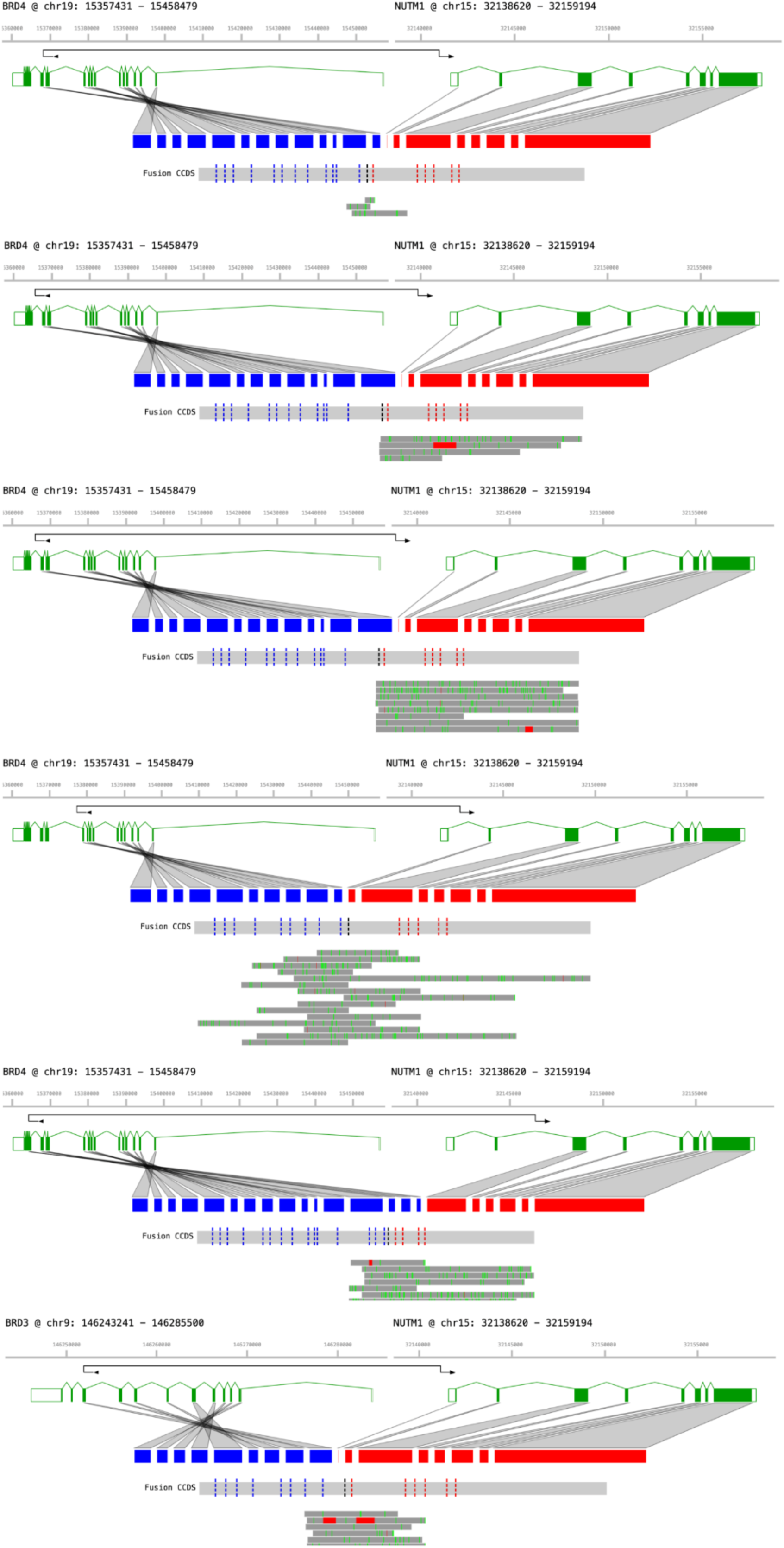

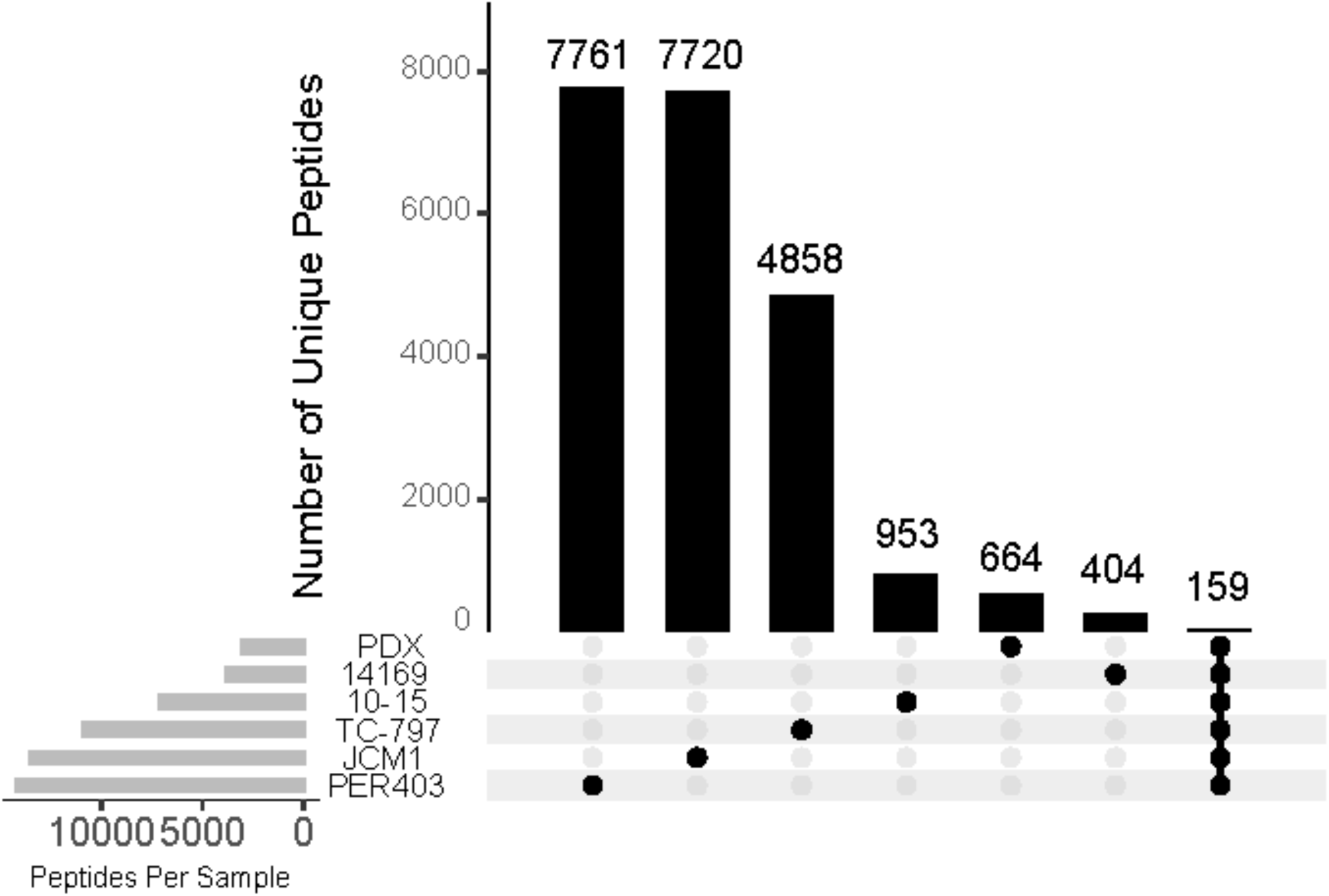

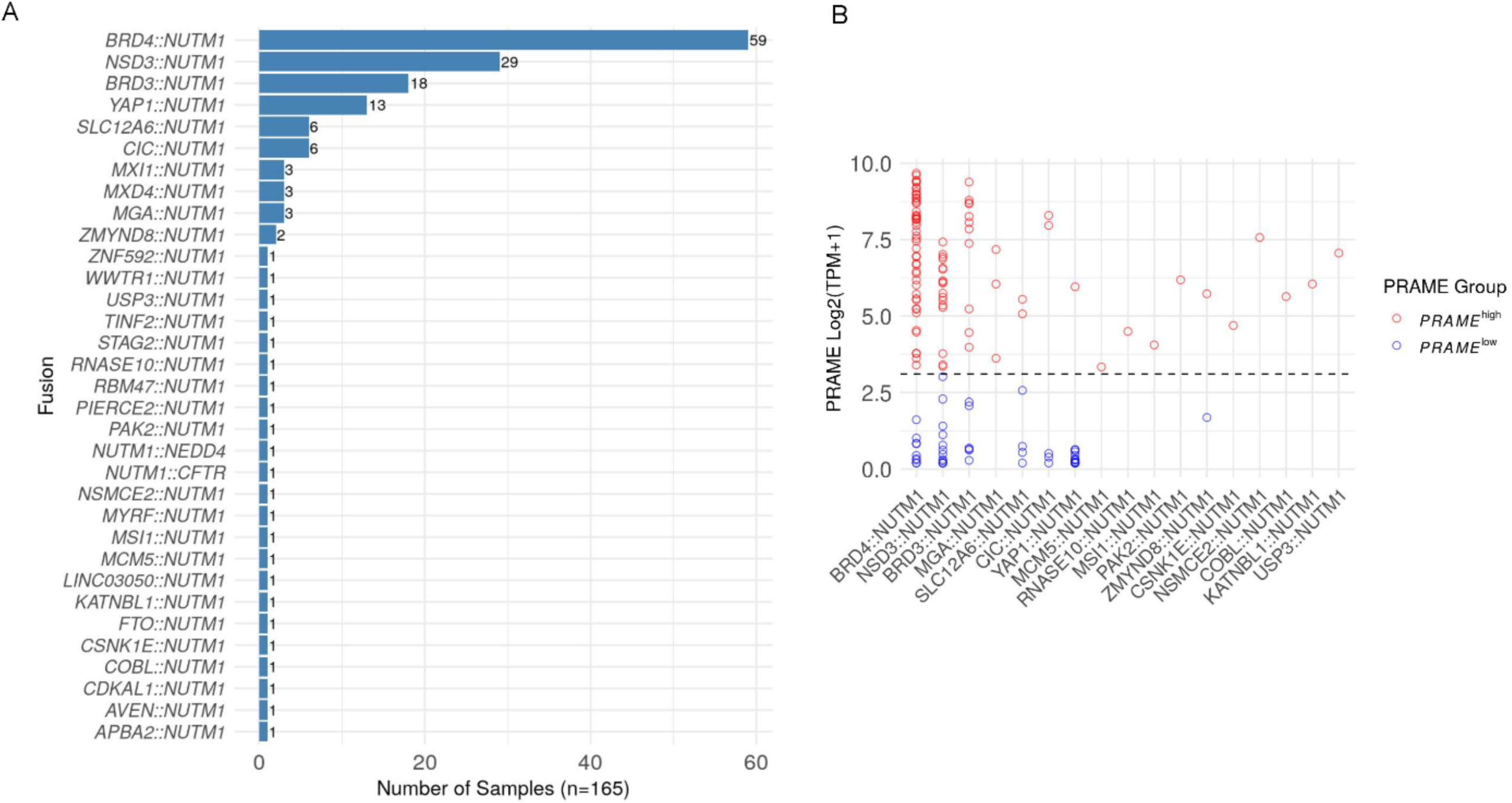

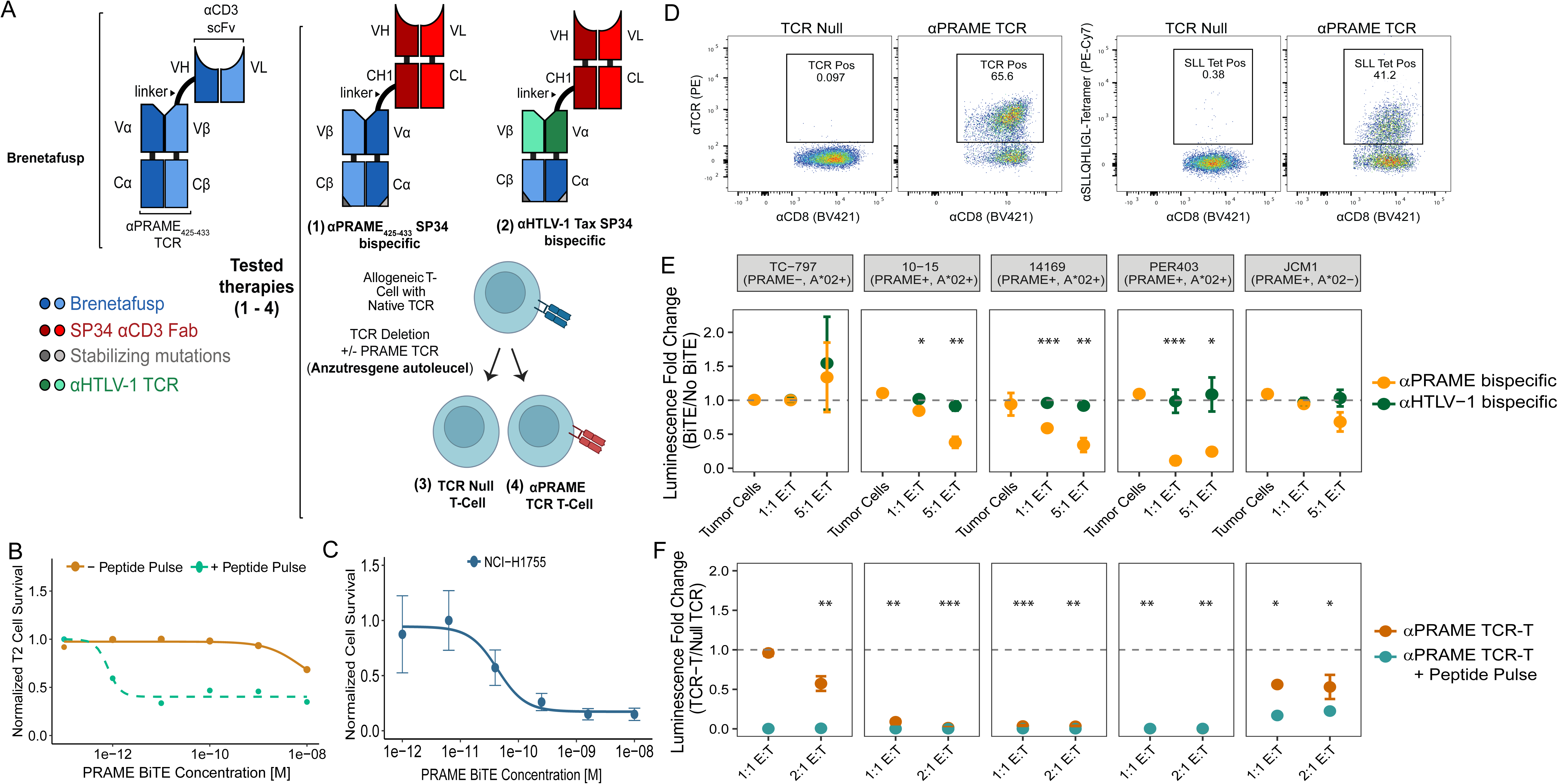

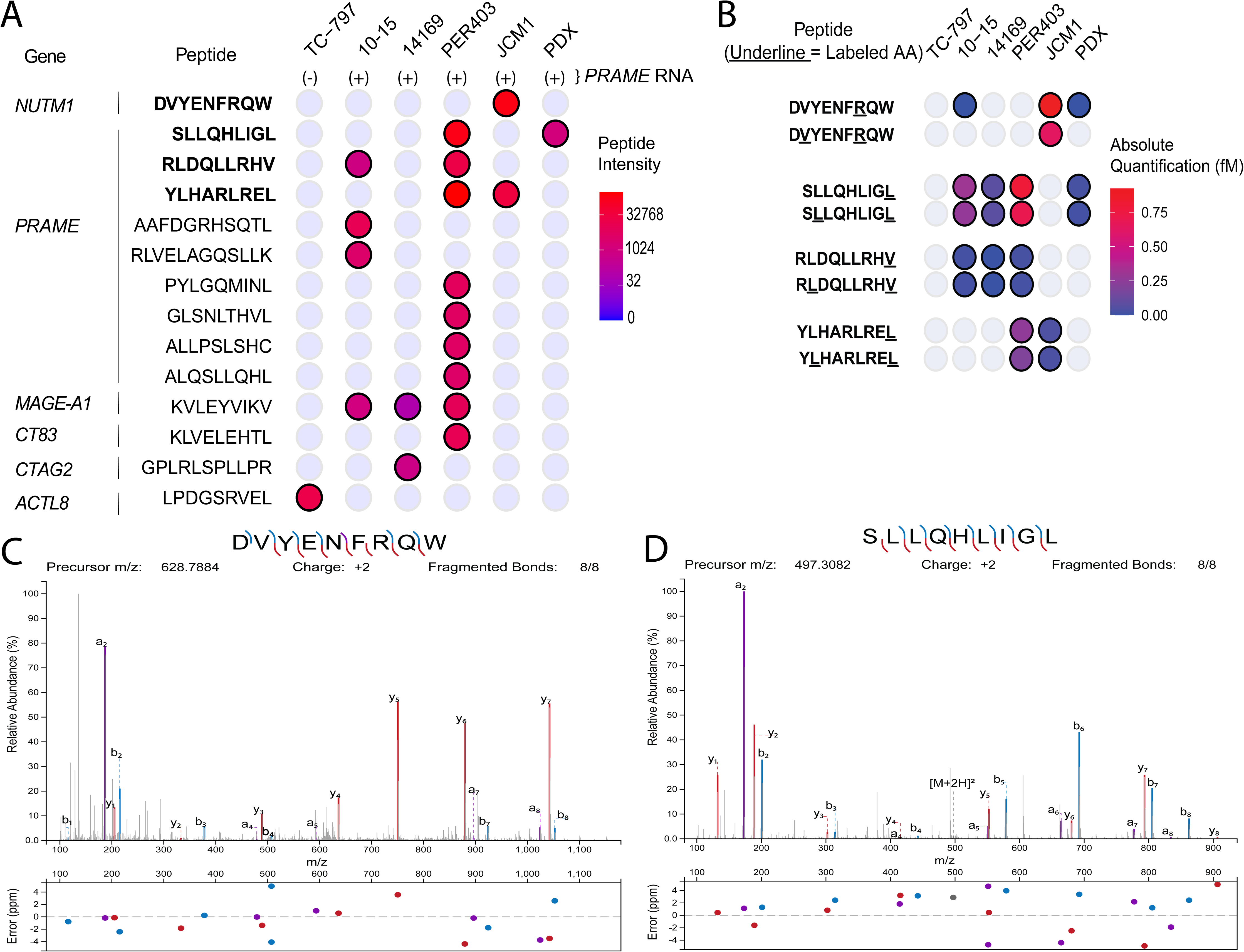

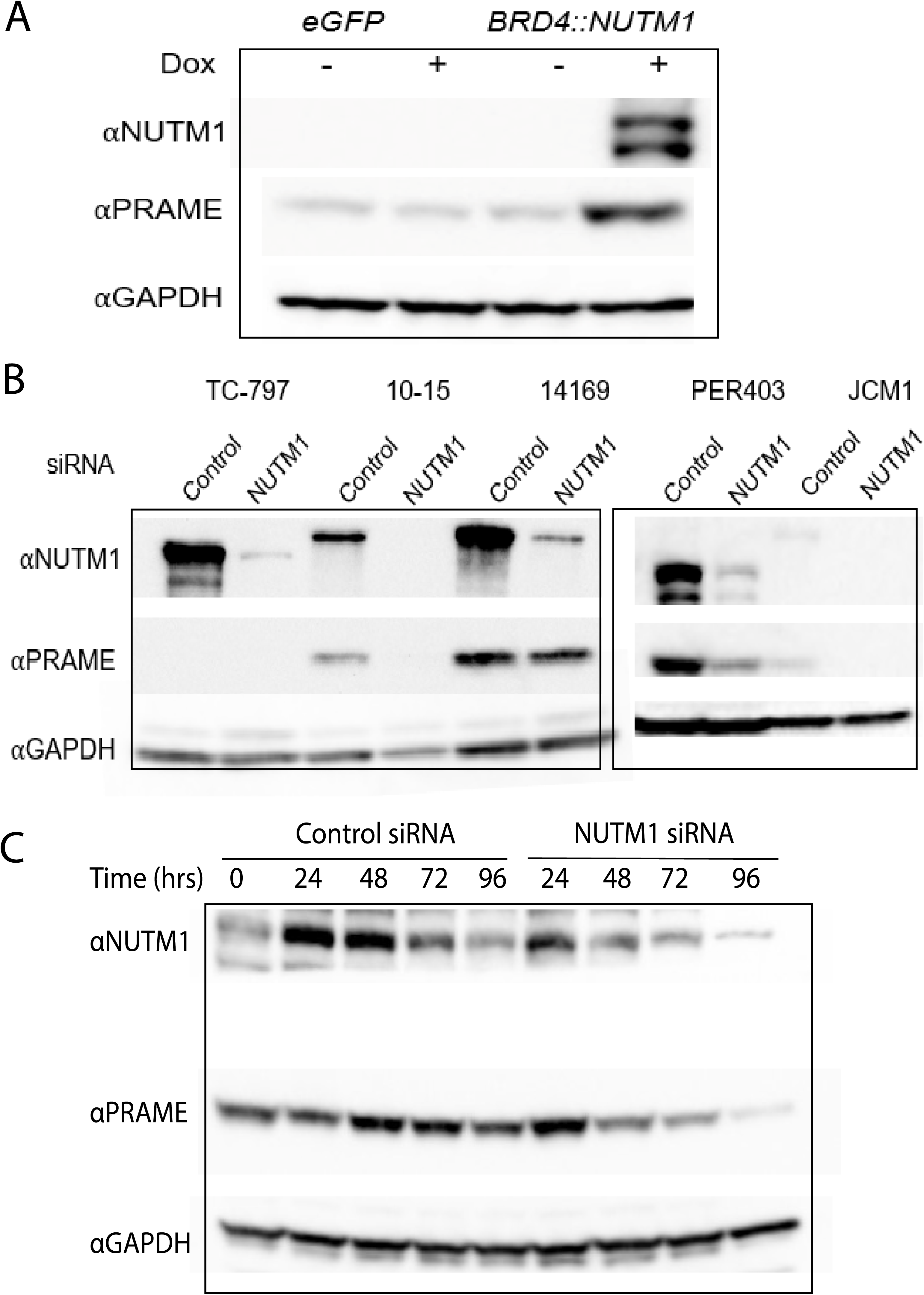

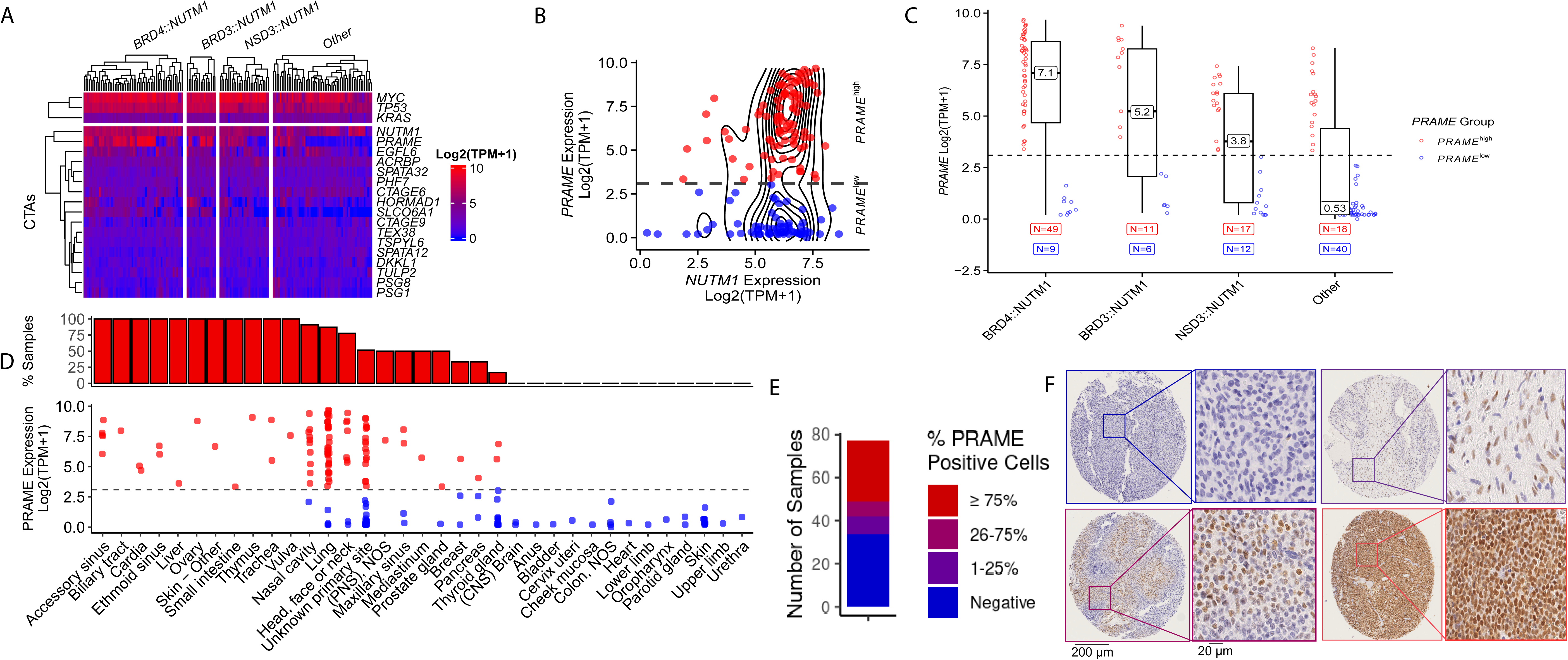

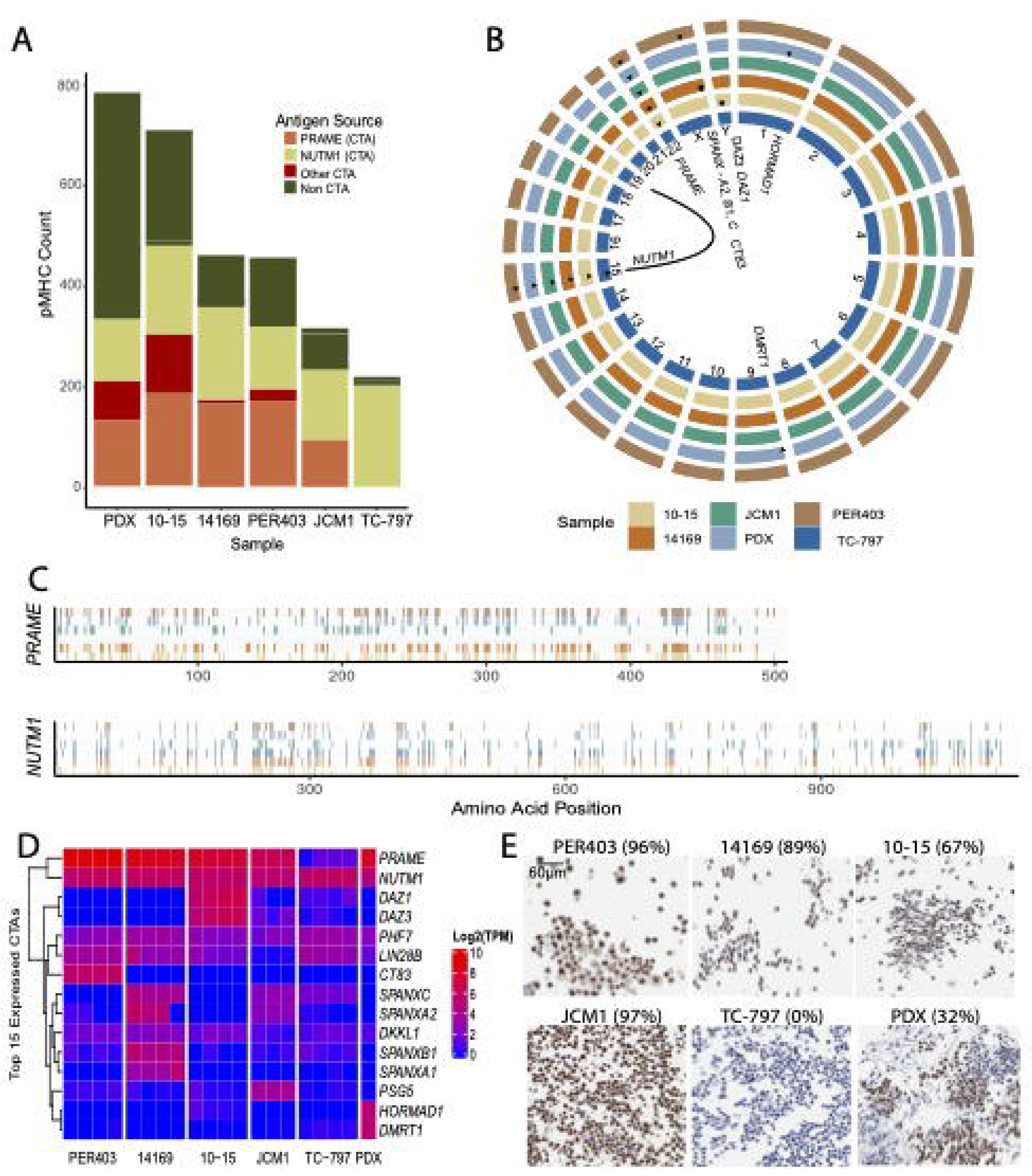

