## Supplementary material for "Shared PRAME Epitopes are T-Cell Targets in NUT Carcinoma": Tables

|  | *HLA-A* | *HLA-B* | *HLA-C* | *PRAME Status* | *Fusion* |
| --- | --- | --- | --- | --- | --- |
| TC-797 | **A*02:01** A*02:01 | B*18:01 B*07:06 | C*05:01 C*07:02 | - | *BRD4::NUTM1* |
| 10-15 | **A*02:01** A*03:01 | B*35:01 B*27:02 | C*03:03 C*02:02 | + | *BRD4::NUTM1* |
| 14169 | **A*02:01** A*29:02 | B*44:02 B*07:02 | C*07:02 C*05:01 | + | *BRD4::NUTM1* |
| PER403 | **A*02:01** A*24:02 | B*08:01 B*35:01 | C*04:01 C*07:01 | + | *BRD4::NUTM1* |
| JCM1 | A*03:01 A*25:01 | B*08:01 B*07:02 |  | + | *BRD4::NUTM1* |
| PDX | **A*02:01** A*01:01 | B*44:02 B*57:01 | C*06:02 C*05:01 | + | *BRD3::NUTM1* |

**Table 1: HLA haplotypes and PRAME status of NUT carcinoma samples.** Bold in HLA-A = HLA-A*02:01+.

| Fusion | Samples (%) | NC Diagnosis (%) | Primary Site | Samples Per Site (%) |
| --- | --- | --- | --- | --- |
| *BRD4::NUTM1* | 59 (36%) | 20 (34%) | Unknown primary site | 15 (25%) |
|  |  |  | Lung | 14 (24%) |
|  |  |  | Nasal cavity | 8 (14%) |
|  |  |  | Accessory sinus | 4 (7%) |
| *BRD3::NUTM1* | 18 (11%) | 6 (33%) | Lung | 7 (39%) |
|  |  |  | Unknown primary site | 4 (22%) |
|  |  |  | Nasal cavity | 3 (17%) |
|  |  |  | Head, face, or neck | 1 (6%) |
| *NSD3::NUTM1* | 29 (18%) | 4 (14%) | Lung | 11 (38%) |
|  |  |  | Thyroid gland | 11 (38%) |
|  |  |  | Head, face, or neck | 2 (7%) |
|  |  |  | Maxillary sinus | 2 (7%) |
| Other | 59 (36%) | 2 (3%) | Unknown primary site | 14 (24%) |
|  |  |  | Lung | 7 (12%) |
|  |  |  | Skin | 7 (12%) |
|  |  |  | Colon, NOS | 4 (7%) |

**Table 2: Cohort characteristics by *NUTM1* gene fusion partner (N=165).**

| **Epitope** | **Single-Labeled Standard** | **Double-Labeled Standard** |
| --- | --- | --- |
| PRAME_312_ RLDQLLRHV | H2N-RLDQLLRH**V***-OH  **V*** = Valine(^13^C_5_, ^15^N) | H2N-R**L***DQLLRH**V***-OH  **L*** = Leucine(^13^C_6_, ^15^N)  **V*** = Valine(^13^C_5_, ^15^N) |
| PRAME_425_ SLLQHLIGL | H2N-SLLQHLIG**L***-OH  **L*** = Leucine(^13^C_6_, ^15^N) | H2N-S**L***LQHLIG**L***-OH  **L*** = Leucine(^13^C_6_, ^15^N) |
| PRAME_462_ YLHARLREL | H2N-YLHARLRE**L***-OH  **L*** = Leucine(^13^C_6_, ^15^N) | H2N-Y**L***HARLRE**L***-OH  **L*** = Leucine(^13^C_6_, ^15^N) |
| NUTM1_237_ DVYENFRQW | H2N-DVYENF**R***QW-OH  **R*** = Arginine(^13^C_6_, ^15^N_4_) | H2N-D**V***YENF**R***QW-OH  **V*** = Valine(^13^C_5_, ^15^N)  **R*** = Arginine(^13^C_6_, ^15^N_4_) |

**Table 3: Single- and double-labeled peptide standards used for targeted mass spectrometry.**
