## Supplementary Tables for "Shared PRAME Epitopes are T-Cell Targets in NUT Carcinoma"

| **Immunoglobulin Target** | **Vendor, Clone (RRID)** | **Dilution** |
| --- | --- | --- |
| *Primary Antibodies* |  |  |
| αNUT | Cell Signaling, Rabbit C52B1 (RRID:AB_2066833) | 1:1,000 |
| αPRAME | Abcam, Rabbit EPR20330 (RRID:AB_3677388) | 1:,2000 |
| α-β-Actin | Sigma-Aldrich, A5441 (RRID:AB_476744) | 1:10,000 |
| *Secondary Antibodies* |  |  |
| αRabbit-HRP | Cell Signaling, Goat #7074S (RRID:AB_2099233) | 1:10,000 |
| αMouse-HRP | Cell Signaling, Mouse #7076S (RRID:AB_330924) | 1:10,000 |

**Supplemental Table 1:** Western blotting antibodies.

| **siRNA Pool** | **Sequences** |
| --- | --- |
| αNUT |  |
| NUTa | AAACUCAGAACUUUAUCCUUA[dT][dT] |
| NUTa antisense | UAAGGAUAAAGUUCUGAGUUU[dT][dT] |
| NUTb | AAACUUCUGUUAGUAAAACAC[dT][dT] |
| NUTb antisense | GUGUUUUACUAACAGAAGUUU[dT][dT] |
| NUTc | AAUUACCUUUGGAAGGAGCUA[dT][dT] |
| NUTc antisense | UAGCUCCUUCCAAAGGUAAUU[dT][dT] |
| NUTd | AAUGGAGAAUCCCAAUGUUGA[dT][dT] |
| NUTd antisense | UCAACAUUGGGAUUCUCCAUU[dT][dT] |
| Scrambled controls |  |
| Scrambled a | GAAAUUACUAUCCACUAUCUA[dT][dT] |
| Scrambled a antisense | UAGAUAGUGGAUAGUAAUUUC[dT][dT] |
| Scrambled b | GAAUUAACACUACUAUGACUA[dT][dT] |
| Scrambled b antisense | UAGUCAUAGUAGUGUUAAUUC[dT][dT] |
| Scrambled c | GAACGAGCUACGGAAUUAUUU[dT][dT] |
| Scrambled c antisense | AAAUAAUUCCGUAGCUCGUUC[dT][dT] |
| Scrambled d | GUAAACAACGUGAGACUUAGU[dT][dT] |
| Scrambled d antisense | ACUAAGUCUCACGUUGUUUAC[dT][dT] |

**Supplemental Table 2:** siRNA sequences.

| NUTM1  CAGE1  PAGE1  PAGE2  PAGE3  XAGE1A  XAGE3  XAGE5  GAGE2A  GAGE12H  GAGE12J  GAGE12E  GAGE13  MAGEA1  MAGEA2  MAGEA4  MAGEA6  MAGEA3  MAGEA9  MAGEA9B  MAGEA10  PSG8  PSG11  PSG1  PSG3  CSH2  PSG5  TRIM64 | MAGEA11  MAGEB1  MAGEB2  MAGEB3  MAGEB5  MAGEB6B  MAGEB16  MAGEB18  MAGEC1  MAGEC2  DAZ1  DAZ3  DAZL  BSPH1  SPAG11A  SPANXA1  SPANXA2  SPANXB1  SPANXC  SPANXD  SPANXN1  PRM1  PRM2  PRM3  AKAP4  TNP2  CGB8  PSG9 | SPANXN2  SPANXN3  SPANXN4  SPANXN5  ODF1  ODF3  ODF4  ROPN1B  SPAM1  SPATA4  SPATA12  SPATA16  SPATA19  SPATA22  SPATA31E1  SPATA31A1  SPATA31A3  SPATA31A5  SPATA31A6  SPATA31A7  SPATA31D3  PRAMEF18  PRAMEF20  CBLL2  CPXCR1  TULP2  EPPIN | SPATA31D4  SPATA32  SPACA1  SPACA3  CATSPERD  CATSPER4  CTAG1A  CTAG1B  CTAGE1  CTAGE6  CTAGE9  SSX1  SSX2  SSX2B  SSX3  SSX4  TSPY1  TSPY2  TSPY3  TSPY4  TSPY8  ACRBP  TSKS  CCDC42  ADAD1  RIMBP3  TENT5D | TSPY9P  TSPY10  TSPYL6  MORC1  TFDP3  LUZP4  DKKL1  SPO11  CRISP2  FMR1NB  TPTE  CT45A2  CT45A7  CT47A1  CT47A2  CT47A3  CT47A4  CT47A5  CT47A6  CT47A7  CT47A11  PRAME  ACTL8  SPEM1  THEG  EGFL6  LIN28B | CT47A12  CT47B1  CT83  TEX13A  TEX19  TEX33  TEX35  TEX37  TEX38  TEX44  TEX101  PATE1  PATE4  PRSS54  PHF7  SLCO6A1  TEKT5  DMRT1  DMRTC2  DNAJB8  SAGE1  LIPI  ADAM2  PLAC1  CABS1  FATE1  HORMAD1 |
| --- | --- | --- | --- | --- | --- |

**Supplemental Table 3:** Manually curated list of CTAs.

| **Construct** | **Amino acid sequence** |
| --- | --- |
| **Primary** | EVQLVESGGGLVQPKGSLKLSCAASGFTFNTYAMNWVRQAPGKGLEWVARIRSKYNNYATYYADSVKDRFTISRDDSQSILYLQMNNLKTEDTAMYYCVRHGNFGNSYVSWFAYWGQGTLVTVSAASTKGPSVFPLAPSSKSTSGGTAALGCLVKDYFPEPVTVSWNSGALTSGVHTFPAVLQSSGLYSLSSVVTVPSSSLGTQTYICNVNHKPSNTKVDKKVEPKSCGGGGSGGGGSGGGGSGGGGSGDAKTTQPNSMESNEEEPVHLPCNHSTISGTDYIHWYRQLPSQGPEYVIHGLTSNVNNRMASLAIAEDRKSSTLILHRATLRDAAVYYCILILGHSRLGNYIATFGKGTKLSVIPYIQNPDPAVYQLRDSKSSDK**F**VCLFTDFDSQ**I**NVSQSKDSDVYITDKCVLDMRSMDFKSNSAVAWSNKSDF**T**CANAFNNSIIPEDTFFPSPESSCGGSGGSGGHHHHHHHH |
| **Anti-CD3 auxiliary** | QAVVTQESALTTSPGETVTLTCRSSTGAVTTSNYANWVQEKPDHLFTGLIGGTNKRAPGVPARFSGSLIGDKAALTITGAQTEDEAIYFCALWYSNLWVFGGGTKLTVLGQPKAAPSVTLFPPSSEELQANKATLVCLISDFYPGAVTVAWKADSSPVKAGVETTTPSKQSNNKYAASSYLSLTPEQWKSHRSYSCQVTHEGSTVEKTVAPTEC |
| **Anti-PRAME auxiliary** | DGGITQSPKYLFRKEGQNVTLSCEQNLNHDAMYWYRQDPGQGLRLIYYSQIMGDEQKGDIAEGYSVSREKKESFPLTVTSAQKNPTAFYLCASSWWTGGASPIRFGPGTRLTVTEDLKNVFPPEVAVFEPS**K**AEIS**R**TQKATLVCLATGFYP**P**HVELSWWVNGKEVH**D**GVCTDPQPLKEQPALNDSRYALSSRLRVSATFWQDPRNHFRCQVQFYGLSENDEWTQDRAKPVTQIVSAEAWGRADC |

**Supplemental Table 4:** Amino acid sequences for primary, anti-CD3 auxiliary, and anti-PRAME auxiliary constructs of PRAME bispecific. CDRs derived from PRAME Vα and Vβ domains are shown in underline. Computationally identified thermostabilizing mutations introduced to PRAME Cα and Cβ domains are shown in bold.
