## Supplementary Figure Legends for "Shared PRAME Epitopes are T-Cell Targets in NUT Carcinoma"

**Supplemental Figure 1** Additional data regarding the Tempus clinical cohort. A) Complete list of *NUTM1* gene fusions represented in clinical cohort. B) *PRAME* expression by gene fusion.

**Supplemental Figure 2** Number of immunopeptides detected in the immunopeptidomes of five NUT carcinoma cell lines and a NUT carcinoma PDX.

**Supplemental Figure 3** Fusion oncogene characterization of NUT carcinoma cell lines and a PDX. From top to bottom: TC-797, 10-15, 14169, PER-403, JCM1, and PDX.

**Supplemental Figure 4** Additional αPRAME BiTE cytotoxicity assay and raw luminescence values. A) First αPRAME bispecific cytotoxicity assay at two time points using luciferase-expressing cell lines. B) Raw luminescent values from the experiment in A. C) Raw luminescent data from the experiment in Figure 5E. D) Raw luminescent data from the experiment in Figure 5F. Error bars = standard deviation.

**Supplemental Figure 5** Complete gating strategy for αTCR and SLLQHLIGL tetramer staining of TCR T-cells.
