## Supplementary material for "Shared PRAME Epitopes are T-Cell Targets in NUT Carcinoma": Figure Legends

**Figure 1** LENS predictions for HLA Class I ligands in NUT carcinoma samples. A) Column plot depicting frequency of different neoantigen types across samples. B) CIRCOS plot describing the CTAs predicted by LENS. Arrows represent CTAs with predicted epitopes. Gene names are given within the circle. Arc represents the *BRD4::NUTM1* fusion (note: PDX has a *BRD3::NUTM1* fusion). C) Predicted epitope landscape within *PRAME* and *NUTM1* with predicted binding affinities of 500 nM or stronger. D) RNA sequencing heat map of cancer/testes genes in these samples. 149 of 164 cancer/testis genes were omitted from the heat map as they were transcriptionally silent. E) PRAME IHC staining on the NUT carcinoma samples with % of PRAME positive cells labeled for each sample.

**Figure 2** *PRAME* RNA and PRAME protein expression in patient samples (N=165) with *NUTM1* fusion oncogenes. A) Heatmap of CTAs by NUTM1 fusion partner. Transcriptionally silent cancer/testis genes (TPM expression less than or equal to 5 TPM in >50% of samples) were omitted (N = 147 / 164). B) *PRAME* expression charted against *NUTM1* expression shows a bimodal distribution. C) *PRAME* expression by *NUTM1* fusion. D) *PRAME* expression by anatomic site. E) Distribution of % PRAME positive cells in a NUT carcinoma TMA. F) Low and high power IHC images from PRAME staining in representative samples (upper left negative, upper right 1-25%, lower left 26-75%, lower right >=75%).

**Figure 3** PRAME levels in NUT carcinoma are enhanced by BRD4::NUTM1. A) Immunoblot of HEK 293T cells with tet-on BRD4::NUTM1 expression showing increased PRAME with BRD4::NUTM1 expression. B) siRNA knockout of BRD4::NUTM1 in NUT carcinoma cell lines using siRNAs results in lower PRAME levels except in *PRAME*- TC-797. Densitometry in 14169 supports reduced PRAME levels with BRD4::NUTM1 loss (not shown). C) PRAME levels are lowest in 14169 four days after transfection with NUT siRNAs.

**Figure 4** PRAME epitopes are presented by HLA Class I molecules on NUT carcinoma cells. A) Summarized results of untargeted immunopeptidomics in NUT carcinoma samples with semi-quantitative peptide intensity shown. B) Summarized results of targeted mass spectrometry with quantitative abundances for selected epitopes. Underlined residues represent isotope-labeled amino acids described in Table 3. C) Representative MS2 spectra of the SLLQHLIGL (PRAME_425_) immunopeptide in PER403 cells. D) Representative MS2 spectra of the DVYENFRQW (NUTM1_237_) immunopeptide in JCM1 cells. Matched fragment ions are labeled.

**Figure 5** NUT carcinoma cells are susceptible to TCR therapeutics modeled after the PRAME BiTE brenetafusp and the TCR T-cell product anzutresgene autoleucel. A) Diagram showing similarities and differences between αPRAME_425_ TCR x SP34 αCD3 BiTE and brenetafusp, as well as the other tested control BiTE and TCR T-cell agents. B) T2 cell peptide pulse data with SLLQHLIGL (PRAME_425_) confirms BiTE specificity for SLLQHLIGL (PRAME_425_). C) Titration of BiTE concentration on the HLA A02+, PRAME+ NSCLC cell line NCI-H1755. D) Flow cytometry with αTCR and SLLQHLIGL tetramer staining confirming ablation of the native TCR and successful transduction with an αPRAME_425_ TCR. E) Cytotoxicity data of the αPRAME_425_ TCR x SP34 αCD3 BiTE and a control αHTLV-1 BiTE against the five NUT carcinoma cell lines. Unpaired t-tests between each condition’s +αPRAME_425_ BiTE and no BiTE were performed and are denoted as follows: * = p<0.05, ** = p<0.01, *** = p<0.001. F) Cytotoxicity data of the αPRAME_425_ TCR T-cells +/- SLLQHLIGL (PRAME_425_) peptide pulse against the five NUT carcinoma cell lines. Similar to above, unpaired t-tests were performed between each αPRAME_425_ TCR T-cell condition and the corresponding TCR null condition with the same significance depiction as in E. Error bars are +/- standard deviation. Raw luminescent values, including additional controls for E&F are provided in Supplementary Figure 4.
